# Microglia coordinate activity-dependent protein synthesis in neurons through metabolic coupling

**DOI:** 10.1101/2025.10.29.685332

**Authors:** Drew Adler, Alejandro Martín-Ávila, Evan Cheng, Mauricio M. Oliveira, Muxian Zhang, Harrison T. Evans, Deliang Yuan, Richard Sam, Nicole D. Zhang, Maria Clara Selles, Olivia Mosto, Wendy J. Liu, Amy X. Guo, Shane A. Liddelow, Robert C. Froemke, Moses V. Chao, Wen-Biao Gan, Eric Klann

**Affiliations:** Center for Neural Science, New York University; New York, 10003, USA; Department of Neuroscience, New York University Grossman School of Medicine; New York, 10016, USA; Institute of Neurological and Psychiatric Disorders, Shenzhen Bay Laboratory; Shenzhen, Guangdong, China; Department of Psychiatry, New York University Grossman School of Medicine; New York, 10015, USA; Department of Otolaryngology, New York University Grossman School of Medicine; New York, 10016; Department of Ophthalmology, New York University Grossman School of Medicine; New York, 10016; Institute for Translational Neuroscience, New York University Grossman School of Medicine; New York, 10016

## Abstract

*De novo* protein synthesis is required for long-lasting synaptic plasticity and memory, but it comes with a great metabolic cost. In the mammalian brain, it remains unclear which cell types and biological mechanisms are critical for sensing and responding to increased metabolic demand. Here we demonstrate that microglia, resident macrophages of the brain, coordinate metabolic coupling between endothelial cells, astrocytes, and neurons to fuel protein synthesis in active neurons. Increasing metabolic demand via a motor task stimulates microglia to secrete the hypoxia-responsive protein CYR61, increasing glucose transporter expression in brain vasculature. Depleting microglia reduces training-induced metabolic fluxes and neuronal protein synthesis, which can be reproduced by blocking CYR61 signaling. Thus, we define a neuroimmune metabolic circuit required for on-demand protein synthesis in mouse motor cortex.

## Main Text

The brain consumes the most energy of any organ in the body by weight and derives ∼95% of its ATP from glucose metabolism ^1^. Of the many energy-intensive processes neurons undergo, protein synthesis is a particular constraint, consuming ∼4 ATP per peptide bond formation ^2^. Indeed, protein synthesis and ATP consumption are tightly correlated ^3^. In neurons, *de novo* protein synthesis provides the substrates required for long-lasting synaptic plasticity, but without energy production via local mitochondria, protein synthesis terminates and dendritic spines fail to grow in response to activating stimuli ^4^. Protein synthesis required for memory consolidation is thought to occur in multiple waves^5^, continuing hours after an initial stimulus. Once potentiated, synapses carry a higher metabolic load, requiring a sustained energetic supply ^6^. Thus, the brain requires a coordinated system to ensure resources are locally allocated to active circuits, both acutely to sustain activity and long-term to maintain plasticity-related processes.

Although neurons can directly use glucose and glycolysis to generate ATP, activity stimulates glucose uptake preferentially into astrocytes, who generate and supply lactate to neurons in a phenomenon known as the astrocyte neuron lactate shuttle (ANLS) ^7–9^. Therefore, neuronal metabolic fueling requires non-cell autonomous communication. Outside of the brain, a core function of tissue resident macrophages is sensing organ level metabolic demand and secreting of factors or phagocytosing products to maintain homeostasis ^10–15^. Microglia, the resident macrophages in the brain, serve several homeostatic functions in adulthood, including regulating synaptic structures ^16–18^, phagocytosing dead cells ^19,20^, and controlling neuronal activity ^21,22^. Recent work in the developing brain suggests depletion of microglia-specific receptors can alter neuronal metabolism^23^. Whether these resident macrophages support ongoing brain metabolism in adulthood, and the mechanism for this support, is presently unknown.

Herein, we use multi-omic approaches, *in vivo* subcellular metabolic imaging, and readouts of behavior-induced *de novo* protein synthesis to demonstrate that microglia actively monitor and maintain local metabolic homeostasis in response to motor activity. Moreover, we propose a core mechanism for microglia-metabolic coordination, the secretion of CYR61 (CCN1), which facilitates the expression of glucose transporter 1 (SLC2A1) in brain endothelial cells: the rate-limiting step of glucose entry into the brain ^24,25^.

### Microglia regulation of brain metabolism in adulthood

To determine whether microglia play an active role in regulating brain energetics broadly, we leveraged an inducible microglia depletion mouse model (Cx3Cr1^CreER^:R26^DTR/+^ hereafter “MgDTR”) in which ∼92% of cortical microglia are depleted four days after commencing diphtheria toxin treatment compared to littermate controls (Cx3Cr1^CreER^:R26^+/+^ hereafter “Control”) given identical treatments (**Figures 1A and 1B**). Re-analyzed proteomic data comparing MgDTR brains to controls using the same model ^16^ identified pyruvate metabolism as the most significantly downregulated set of proteins in MgDTR mice (**Figure S1A**).

**Fig. 1.**
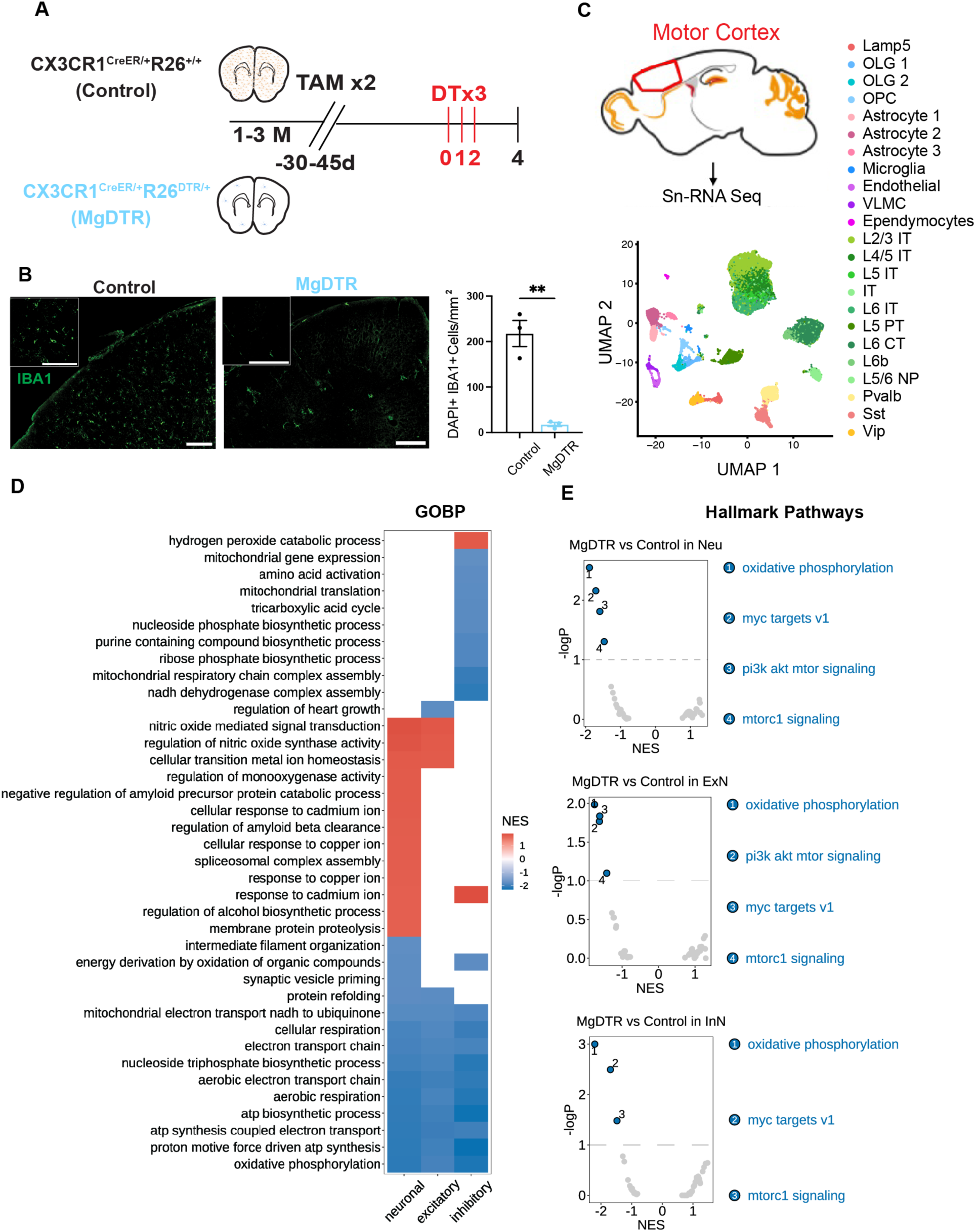
Microglia depletion leads to brain metabolic dysregulation. (**A**) Experimental time course with representative images (**B**) with zoomed insets and quantification of inducible microglia depletion with diphtheria toxin (N_Cntr_=3, N_MgDTR_=3). (**C**) (top) approximate extracted region for sn-RNA Seq and (bottom) UMAP representation of 55,466 nuclei from MgDTR and control mice. Each dot represents a different nucleus, and each color represents a predicted cluster (N_Cntr_=3, N_mgDTR_=3). (**D**) GSEA on all “GOBP” from select neuronal clusters or aggregate neuronal clusters with normalized enrichment score (NES) heatmap. Colored boxes indicate FDR q<.1. (**E**) GSEA on all “Hallmark Gene Sets” from neuron, excitatory neuron, and inhibitory neuron aggregate clusters. Two-tailed unpaired t-test for (B) **P<.01. Scale bar: 200µm for (B), Illustration in (A) created with BioRender.com. Neu= neuron, ExN= excitatory neuron, InN= inhibitory neuron.

To identify candidate cell types with altered metabolism as a consequence of microglia depletion, we conducted single-nucleus RNA sequencing (snRNA-seq) with unbiased clustering on cells from the motor cortex of MgDTR mice, a region where homeostatic roles for microglia have been demonstrated previously ^16,26^. We generated an atlas of 55,466 nuclei and represented the data using dimensionality reduction by use of uniform manifold approximation and projection (UMAP). Unbiased clustering using Seurat predicted 23 nuclei clusters containing neurons, glia, and stromal cells (**Figures 1C, S1B, and S1F-S1I**). As expected, canonical microglia genes including *P2ry12*, *Hexb*, and *Siglech* were significantly reduced in MgDTR mice after pseudobulking RNA from all cells in each group (**Figures S1C-S1E**). Gene set enrichment analysis (GSEA) on nuclei clusters identified multiple metabolic gene sets that were dysregulated in neuronal, glial, and vascular populations from MgDTR mice, including oxidative phosphorylation, glycolysis, aerobic electron transport chain, and ATP biosynthetic process (**Figure 1D, 1E, S1J, S1K, and Data S1 to S2**). We found that 79% (11/14) of the unenriched gene ontology biological pathways (GOBP) discovered in an aggregate neuronal population (FDR q value<.1) were directly related to ATP synthesis (**Figures 1D and Data S1**). Surprisingly, neuronal populations showed broad decreases in ATP synthetic pathways as well as the canonical mTORC1 pathway, one of the major signaling cascades that triggers *de novo* protein synthesis. In contrast, glia and vascular cells showed broad increases in ATP synthetic pathways (**Figure S1J and S1K**).

Changes to metabolic pathways at the transcriptional level do not necessarily demonstrate direct changes in intracellular metabolic fluxes, as they may reflect compensations to inadequate precursors. To more specifically characterize the relevance of our transcriptomic findings we used a new intracellular ATP sensor, iATPSnFR2 ^27^, and monitored real-time *in vivo* ATP fluxes in motor cortical neurons of MgDTR mice and non-depleted control littermates during a forced treadmill running paradigm to increase local metabolic demand. We first confirmed that our paradigm increased neuronal activity by measuring Ca^2+^ in L2/3 somas and L1 dendrites using Thy1.2 gCaMP6s mice (**Figures S2A-S2G**). To measure real-time intracellular fluxes of ATP in neurons, we injected the motor cortex with adeno-associated viruses (AAVs) packaging iATPSnFR2-mCherry under the neuronal promotor hSyn1 and imaged L2/3 somas and L1 neuropil during short (100 frame, 3.6 min) and longer (250 frame, 9 min) runs (**Figure 2A**). As a positive control to test the specificity of the sensor, we blocked mitochondrial ATP production in anesthetized mice using the F-ATPase inhibitor oligomycin A, which led to a decrease in iATPSnFR2 fluorescence in L1 neuropil (**Figure S2H**). Previous studies evaluating activity-dependent ATP fluxes in neurons during physiological activity *in vivo* suggest ATP levels are either invariant during activity ^28^ or broadly increase ^29^, but neither of these studies reported ATP levels in subcellular compartments. Our results suggest that in mice with functional microglia, ATP in L1 neuropil dips below baseline during running, and during extended activity, slightly exceeds baseline in the post-running state (**Figures 2B and 2C**). Conversely, ATP increases in L2/3 somas after commencing motor training and remains elevated in the post-run resting state, perhaps as a predictive allocation of future demand ^29^ (**Figures 2G and 2H**). These compartment-specific differences may reflect higher metabolic demands in neuronal processes than soma ^6^. During 2-3 short runs, compartment-specific ATP levels were unchanged in MgDTR mice. However, after extended runs ATP fluxes were significantly reduced in both L1 (**Figure 2D and 2E**) and L2/3 (**Figure 2H and 2J**) in the post-run recovery phase, ∼26 mins after the commencement of running. These results suggest microglial input is required for continual ATP maintenance after extended periods of activity.

**Figure 2.**
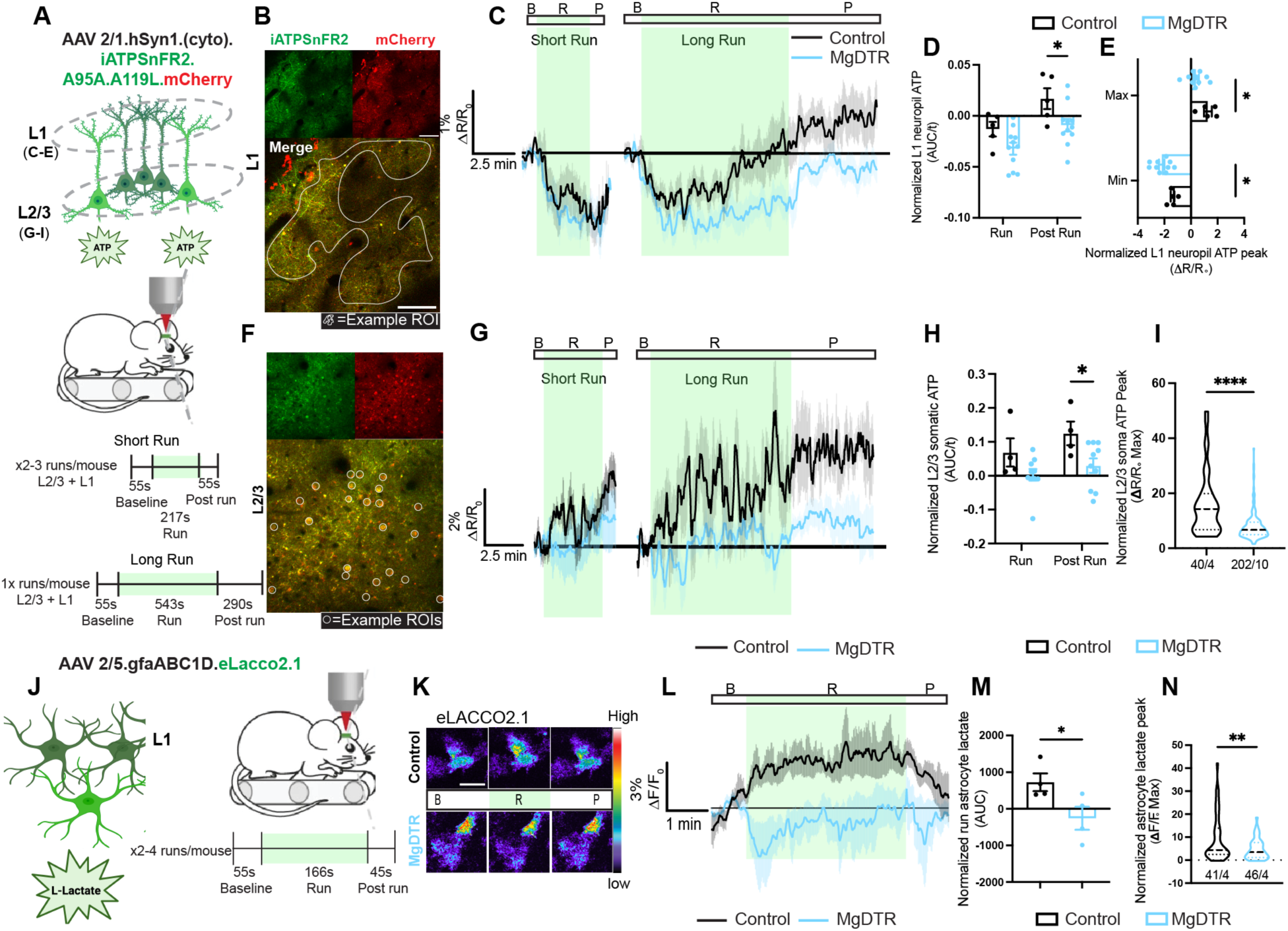
Activity-dependent metabolic fluxes in neurons and astrocytes require microglia. (**A**) Schematic of iATPSnFR2-mCherry ATP sensor in neurons and experimental design for imaging ATP flux in motor cortical neurons during treadmill running. (**B**) Representative image of iATPSnFR2-mCherry signal in L1 motor cortical neuropil. (**C**) Mean animal 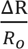 L1 neuropil ATP traces during baseline (B), run (R), and post run (P) phases of short and long runs for control (black) and MgDTR (blue) mice. (N_Cntr_=5 mice, N_MgDTR_=11 mice). (**D**) Integrated iATPSnFR2 signal during run and post run phases per mouse normalized by phase time (t). (**E**) maximum and minimum ATP signal per ROI (N_Cntr_=1 ROI/mouse from 5 mice, N_MgDTR_=1 ROI/mouse from 11 mice). (**F**) Representative image of iATPSnFR2-mCherry signal in L2/3 motor cortex. (**G)** Similar to (C) but for mean animal L2/3 somatic iATPSnFR2 signal (N_Cntr_= 4 mice, N_MgDTR_=10 mice). (**H**) Similar to (D) but for mean mouse L2/3 somatic signal (N_Cntr_=4 mice, N_MgDTR_=10 mice). (**I**) Peak somatic 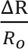 per neuron (N_Cntr=_= 40 cells from 4 mice, N_MgDTR_= 202 cells from 10 mice). (**J**) Schematic illustration of eLACCO2.1 extracellular lactate sensor in astrocytes and experimental design. (**K**) representative ROI heatmaps of control and MgDTR eLACCO2.1 signal and (**L**) mean animal 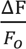 traces during running phases (N=4 mice each genotype). (**M**) Integrated extracellular lactate signal during the run phase per mouse and (**N**) maximum 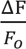 signal per astrocyte (N_cntr_=41 cells/4 mice, N_MgDTR_=46 cells/4 mice). Shading represents +/- S.E.M. two-tailed unpaired t test for (I), (M), and (N); Repeated measures mixed effects analysis with Sidak’s post hoc test for (D), (E), and (H); *P<.05, **P<.01, ****P<.0001. Scale bar: 75 µm for (B) and 50 µm for (K). Illustrations (A), and (J) created with BioRender.com.

Considering that energetic pathway changes were also evident in glia and stromal cells, we asked whether ATP deficits in neurons reflected inadequate metabolic shuttling. In gray matter, oxidized lactate, which has been proposed to fuel activity-dependent ATP synthesis in neurons, is thought to be derived from astrocytes, which convert blood glucose to lactate in the ANLS ^8,30^. To determine whether microglia depletion altered lactate release from astrocytes, we expressed the extracellular lactate sensor eLACCO2.1 ^31^ in motor cortical astrocytes using AAV 2/5.gfaABC1d-eLACCO2.1 in MgDTR mice and controls and imaged L1 during treadmill running (**Figure 2I**). We validated the sensitivity of eLACCO2.1 to extracellular lactate *in vivo* by applying L-lactate directly to an open cranial window while blocking the import of lactate with monocarboxylate transporter1/2 (MCT1/MCT2) inhibitor AR-C155858 ^32^, which increased the fluorescent signal (**Figures S2I-S2K**). In control mice lactate levels consistently increased around astrocytic processes during motor training. However, in MgDTR mice, mean extracellular lactate levels failed to increase above baseline and were significantly lower than controls (**Figures 2J-2M**). Together, these results suggest neuronal activity-dependent ATP deficits in the absence of microglia may reflect a breakdown of the ANLS.

### Microglia regulation of activity-dependent protein synthesis in neurons

Given the requirement of ongoing oxidative metabolism for neuronal protein synthesis ^4^, the specific involvement of astrocytic lactate in plasticity- and learning-associated protein synthesis ^33,34^, and the role of microglia in motor plasticity ^16^, we tested the hypothesis that microglia regulate neuronal protein synthesis.

Although motor skill learning is known to be protein synthesis-dependent ^35,36^, changes in *de novo* protein synthesis following a motor task have not been reported. Therefore, to determine levels of protein synthesis induced by motor training in wild-type (WT) mice we used our recently developed method for tracking behaviorally linked *de novo* translation *in vivo* ^37^. This approach employs retro-orbital delivery of the methionine analogue L-azidohomoalanine (AHA) in awake mice, enabling the measurement of *de novo* protein synthesis via fluorescent non-canonical amino acid tagging (FUNCAT) ^38^. We delivered AHA and then ran mice on a rotarod set to a constant speed for 1 hour to mimic activity evoked in our treadmill-running paradigm without the need for surgical head bar placement. 30 mins after training we evaluated labeling of newly synthesized proteins (**Figure 3A**). Compared to AHA-injected mice that were returned to their home cage (HC), running mice exhibited 2-3–fold increases in L2/3 and L5 *de novo* protein synthesis of the motor cortex (**Figures 2B-2D**). Furthermore, the effect appeared to be region-specific as levels of protein synthesis were significantly greater in the motor cortex than the nearby somatosensory cortex (**Figure 2E**). FUNCAT labeling in CamKII^Cre/+^;R26^tdtomato^ mice confirmed that this increase occurred predominantly in excitatory neurons (**Figure S3**).

**Figure 3.**
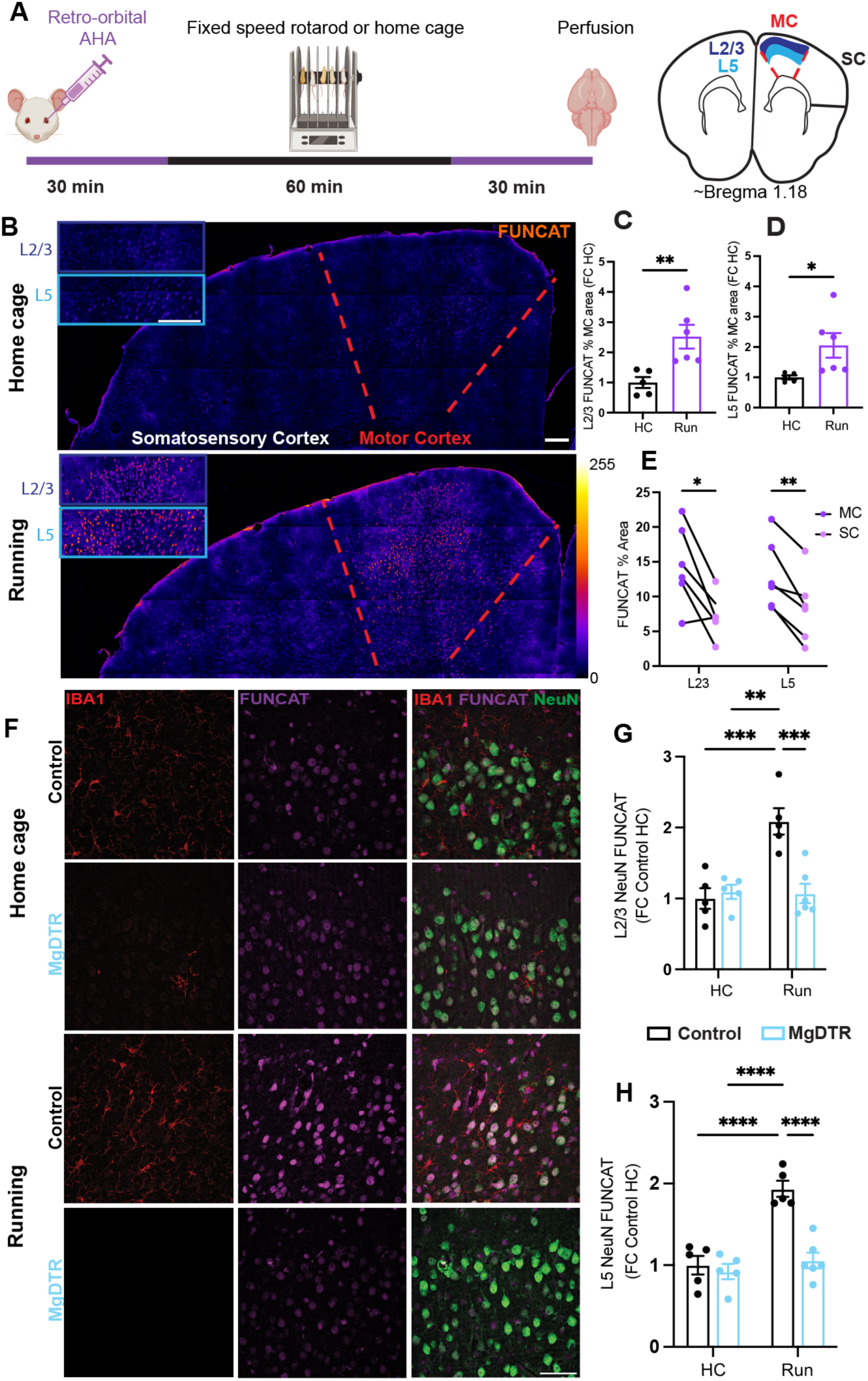
Motor task-induced *de novo* protein synthesis requires microglia. (**A**) Schematic of “*In vivo* FUNCAT” used with rotarod training and illustration of approximate coronal slice used for analysis. (**B**) Representative sections with FUNCAT heatmaps and zoomed L2/3 (dark blue frame) and L5 (light blue frame) insets for HC and Running groups. Red dotted area indicates approximate motor cortex. Quantification of % MC area in (**C**) L2/3 and (**D**) L5 with FUNCAT positivity normalized to mean HC value (N_HC_=5 mice, N_Run_=6 mice). (**E**) % FUNCAT area in MC vs SC within Running mice (N_Run_=6 mice). (**F**) Representative images of microglia (IBA1+, red), FUNCAT (purple), and merged signal with neurons (NeuN+, green) from L2/3 MC Running or Home cage MgDTR or control mice. Quantification of relative neuron FUNCAT signal per animal normalized to mean Control HC value for (**G)** L2/3 or (**H**) L5 (N_CHC_=5 mice, N_MHC_=5 mice, N_CRun_=5 mice, N_MRun_=6 mice). Unpaired two-tailed *t* test for (C) and (D); repeated measures mixed effects model with Sidak’s post hoc test for (E); two-way ANOVA with Tukey’s post hoc test for (G) and (H). Genotype x Condition interaction for (G) and (H) respectively F(1,17)=14.62, 15.45, **P=.0014, .0011. *P<.05, **P<.01,***P<.001 ****P<.0001. Scale bar: 200µm for (B), 50 µm for (F). (A) (left) Illustration created with BioRender.com. MC= motor cortex, SC= somatosensory cortex.

To determine whether neuronal protein synthesis requires homeostatic input from microglia, we repeated our motor FUNCAT paradigm in MgDTR mice. In resting mice, microglia depletion had no effect on basal levels of protein synthesis. However, in the running cohort, microglia depletion abrogated increases in neuronal protein synthesis in both L2/3 (**Figures 3F and 3G**) and L5 (**Figure 3H**). Thus, motor training-induced increases in motor cortical neuronal protein synthesis requires functional microglia.

We hypothesized that the deficits in training-induced *de novo* translation in neurons from MgDTR mice was related to deficits in the ANLS and neuronal ATP synthesis demonstrated in our imaging studies (**Figures 1G-1S**). Although the stimulation of astrocytic glycolysis and lactate release by activity is well supported ^39–41^, whether lactate is a primary source for activity-dependent neuronal ATP production remains controversial ^42,43^. We directly tested the involvement of lactate for the generation of neuronal ATP fluxes during physiological behavior by comparing L1 ATP fluxes before and after cranial window application of α-cyano-4-hydoxcinnamic acid (4-CIN), a potent blocker of the neuron specific MCT2 lactate transporter ^44,45^ in wild-type mice (**Figures 4A and 4B**). Phenocopying ATP fluxes in MgDTR mice, application of 4-CIN had no effect during short runs, but during long runs it blocked the recovery and post-run excess of ATP (**Figures 4C-4E**). This suggests lactate is not immediately required for neuronal ATP synthesis but is required during periods of extended neuronal activity.

**Figure 4.**
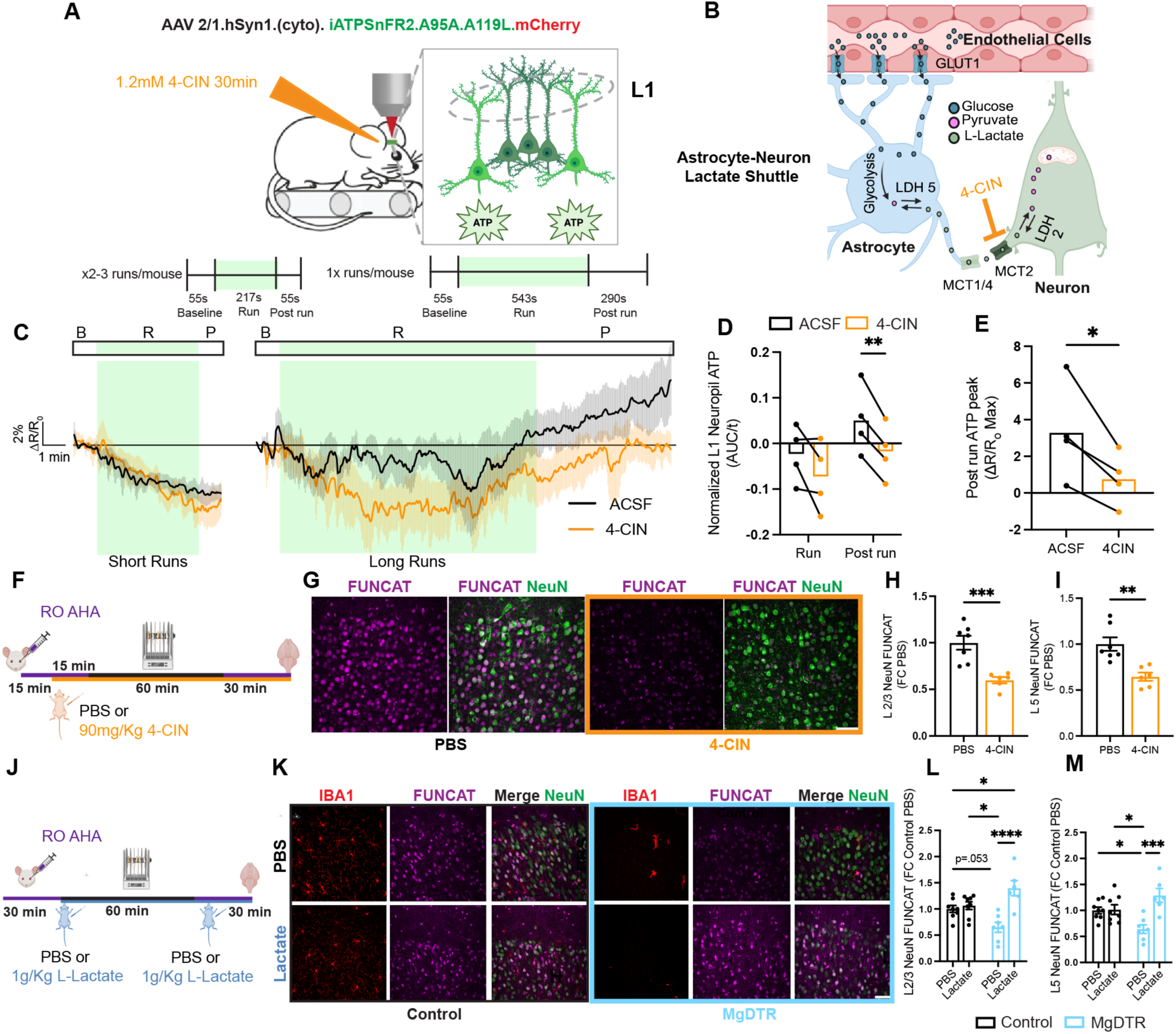
Microglia regulate training-induced *de novo* protein synthesis through metabolic support. (**A**) Schematic of iATPSnFR2 sensing in L1 neuropil with application of 4-CIN. (**B)** Illustration of the ANLS^43^ and blockade of lactate transporter MCT2 with 4-CIN. (**C**) Mean animal 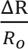 L1 neuropil ATP traces during before (ACSF, black) and after drug application (4-CIN, orange) during short and long runs (N=4 mice, each given both treatments). Shading represents S.E.M. (**D**) Integrated iATPSnFR2 signal during run and post run phases per mouse normalized by phase time (t). (**E**) Peak ATP signal per ROI (1 ROI per animal/4 animals per condition). (**F to I**) (F) Schematic illustration of experimental setup, (G) L2/3 representative images, and quantification of (H) L2/3 and (I) L5 neuronal (green) FUNCAT signal (purple) in mice injected with saline (black) or 4-CIN 15 min prior to motor training (N_PBS_=7 mice, N_4CIN_=6 mice). FUNCAT positivity normalized to mean PBS value. (**J** to **M)** (J) Schematic illustration of experimental setup, (K) L2/3 representative images with microglia (IBA1+, red), neurons (NeuN+, green),and FUNCAT (purple); and quantification of (L) L2/3 and (M) L5 neuronal FUNCAT signal in Control (black) or MgDTR (light blue) mice injected with saline or L-lactate immediately prior to and after motor training normalized to Control HC mean (N_CPBS_=8 mice, N_CLac_=8 mice, N_MPBS_=7 mice, N_MLac_=6 mice). Repeated measures mixed effects analysis with Sidak’s post hoc test (D); Paired two-tailed *t*-test for (E); Unpaired two tailed *t*-test for (H) and (I); Two-way ANOVA with Tukey’s post hoc test for (L) and (M). Genotype x Condition interaction for (L) and (M) respectively F(1,25)=13.4, 11.69 15.45, **P=.0012, .0022. *P<.05, *P<.05, **P<.01, ***P<.001 ****P<.0001. Scale bar: 50 µm for all images. (A, right), (B), (F), and (J) illustrations created with BioRender.com.

Deficits in activity-dependent ATP synthesis quickly lead to synaptic dysfunction ^46^ as well as deficits in protein synthesis and plasticity ^4^. We asked whether blocking lactate transport in neurons was sufficient to block motor task-induced increases in neuronal protein synthesis in the motor cortex. To address this question, we injected 4-CIN, which is known to permeate the blood-brain barrier rapidly^47^, into wild-type mice 15 mins prior to the rotarod task and compared the neuronal FUNCAT signal to saline-injected controls subjected to the same paradigm (**Figure 4F**). Mice injected with 4-CIN prior to the motor task had a 35-40% reduction in neuronal FUNCAT labeling in L2/3 and L5 compared to controls, similar to our results in MgDTR mice (**Figures 4G-4I**). These results suggest neuronal lactate uptake is necessary for activity-dependent protein synthesis in the motor cortex.

Finally, we tested whether deficits in neuronal protein synthesis in MgDTR mice could be rescued by supplying exogenous lactate. We injected MgDTR mice and their littermates with either saline or L-lactate immediately prior to and after the rotarod task finished and compared the FUNCAT signal in motor cortical neurons 30 mins later (**Fig. 3J**). In controls, injection of lactate did not further increase *de novo* protein synthesis in motor cortical neurons. However, in MgDTR mice, lactate injection completely rescued deficits in neuronal *de novo* protein synthesis, and in L2/3, even slightly exceeding the FUNCAT signal of saline-injected control mice (**Figures 4K-4M**). Together these results suggest microglia regulate protein synthesis in neurons via modulation of brain lactate transport, which is required for neuronal ATP synthesis during periods of sustained demand.

### Motor training activates microglia metabolic coupling

Microglia are known to sense and respond to brain activity through increased communication with neurons ^21,22,48–50^, astrocytes ^51–53^, and endothelial cells ^54–56^. To determine the impact of motor activity on microglia, we first evaluated whether microglia change morphologically in response to motor training. Motor cortical microglia from running mice exhibited an increase in somatic size and a decrease in ramification, consistent with an activated phenotype ^57^ (**Figures 5A-5G**). Next, we stained for the microglia structural marker IBA1 in FUNCAT-labeled slices from motor trained or HC mice to determine whether microglia themselves respond to training by increasing *de novo* protein synthesis. Indeed, microglia in the motor cortex of trained mice exhibited an increase in *de novo* protein synthesis (**Figures 5H and 5I**).

**Figure 5.**
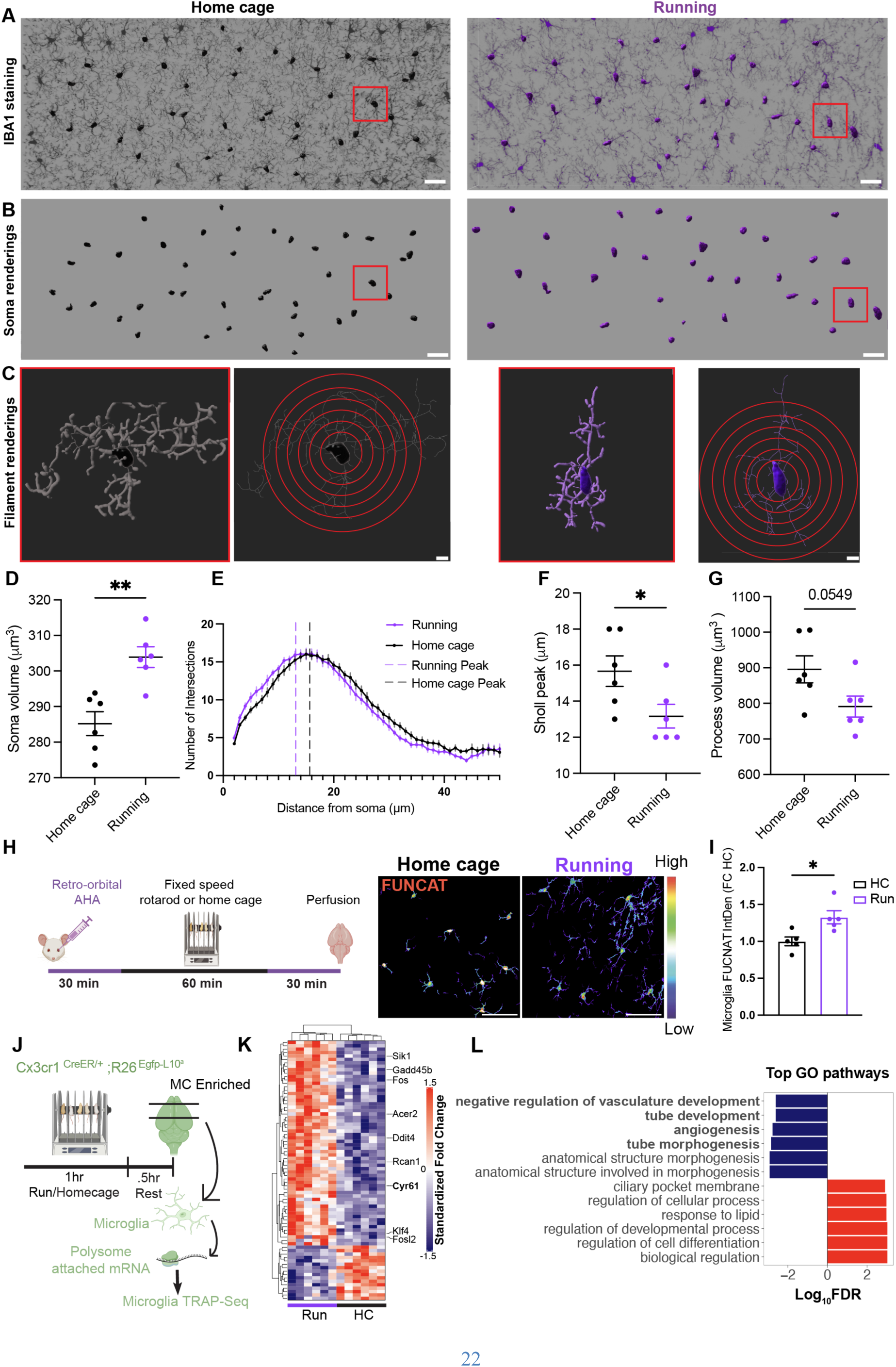
Microglia respond to motor training through morphological and transcriptomic changes. (**A**) Representative L2/3 microglia (IBA1+) in home cage control (black) and running mice (purple). Red box indicates the representative microglia depicted in (C). (**B**) Imaris representative soma renderings. (**C**) Right, Imaris “filament” renderings depicting microglial processes from two representative microglia shown in red boxes above. Left, the same microglia skeletonized with superimposed schematic of Sholl analysis. (**D**) Mean microglial soma volume per mouse (N= 6 mice per group, n>38 microglia/slice, n>127 microglia/animal). (**E**) Mean group Sholl analysis from home cage and running mice. Dotted lines represent the mean Sholl peak per group defined as the distance from the soma with the maximum number of intersections. (**F**) Mean microglial Sholl peak per mouse. (**G**) Mean microglial process volume per mouse. (**H**) Schematic of experiment and FUNCAT heatmap within IBA1+ microglia masks in HC (black) and Running (purple) mice. (**I**) respective quantification of mean integrated density (IntDen) of FUNCAT signal normalized to mean HC value per analysis (N_HC_=5 mice, N_Run_=5 mice). (**J**) Schematic of microglia TRAP-seq: mRNA bound to ribosomes from microglia enriched from the motor cortex of Cx3cr1^CreERT2/+^:R26^EGFP-L10a/+^ mice were sequenced in Running mice trained for 1 hour and rested .5hrs or HC littermates (N= 6 per genotype). (**K**) Hierarchical clustering and heatmap of fold changes for all differentially expressed transcripts (P_adjusted_<.05, 77 total) in individual Running (purple) and HC (black) replicates. (**L**) GO analysis top 6 up (red) and down (blue) pathways discovered in microglia TRAP-seq. Unpaired two-tailed student’s *t* test (D), (F), (G), and (I). *P<.05, **P<.01. Scale bar: 30 µm for (A) and (B), 5 µm for (C), and 50 µm for (H). Illustrations in (H) and (J), created with BioRender.com.

To identify the specific mRNA transcripts that are translated in microglia after training that might regulate the ANLS, we generated microglia-translating ribosome affinity purification (TRAP) ^58^ mice (Cx3cr1^CreER^R26^EGFP-L10a/+^) and compared the ribosome-bound mRNAs in microglia between running and HC mice (**Figure 5J**). Principal component analysis and hierarchical clustering revealed distinct clustering of microglia from running and HC mice (**Figures 5K and S4A**). Validating our TRAP pulldowns successfully targeted microglia, we detected an enrichment in microglial specific genes (*Cx3cr1*, *C1qa*, and *C1qb*) in the TRAP fraction compared to the total lysate fraction, which contains RNA from all cells and was enriched in non-microglial genes (**Figure S5B**). We identified 77 differentially regulated transcripts (**Figures 5C, 5D, S4C, and Data S5;** 60 up, 17 down, P_adjusted_<.05). Supporting the FUNCAT results in microglia, GSEA ^59^ on the TRAP-seq dataset revealed the top gene ontology (GO) pathway increased in running mice was “cytosolic ribosome” along with significantly enriched pathways (nominal P value <.01) “translation at synapse”, “cytosolic large ribosomal subunit”, “cytosolic small ribosomal subunit”, and “cytoplasmic translation” (**Figure S4E**). In addition, we noticed a strong overlap between upregulated transcripts in microglia from running mice and of transcripts from Mecp2-null macrophages, which have increased transcription of glucocorticoid and hypoxia-responsive genes ^60^. Indeed 27% of upregulated ribosome-bound mRNAs in microglia from running mice were identified in at least 1 hypoxia gene set ^60^ suggesting that a key metabolic signal that microglia may sense is local changes in oxygen (**Figure S4D**). We then focused on pathways and upregulated transcripts that might be involved in microglia-astrocyte interaction, which in turn could regulate the function of the ANLS, such as secreted cytokines or ephrin kinases ^61,62^. Unexpectedly, microglia-endothelial interaction emerged as a more compelling nexus, as many of the most significant differentially regulated GO pathways were related to endothelial function such as “tube morphogenesis”, “angiogenesis”, and “negative regulation of vasculature development” (**Figure 5L**).

Microglia have emerged as a key regulator of brain endothelial function in homeostatic conditions ^54–56^. We asked whether there were genes in the vascular cell cluster from our MgDTR Sn-RNAseq dataset that might corroborate the metabolic dysfunctions found in the MgDTR mice. Indeed, *Glut1* (*Slc2a1)*, the primary endothelial glucose transporter facilitating glucose entry from blood into the brain ^63^, was one of the most downregulated genes (Log_2_FC= -.7146, P_adjusted_= 9.2276E-12) in the vascular cluster (**Figure 6A**). Furthermore, GLUT1 protein is known to increase in motor cortical endothelial cells after running ^64^, providing a mechanism for local metabolic facilitation. To determine whether microglia regulate activity-dependent brain endothelial GLUT1 expression, we measured GLUT1 protein in tomato lectin+ endothelial cells in running and HC MgDTR mice with immunofluorescence. As expected, motor training increased motor cortical endothelial GLUT1 in control mice, which was absent in the MgDTR mice (**Figures 6B and 6C**). Furthermore, there was a significant main effect of genotype (microglia presence) on endothelial GLUT1 protein levels (P=.0279). Together, these findings suggest that microglia actively regulate the expression of endothelial GLUT1.

**Figure 6.**
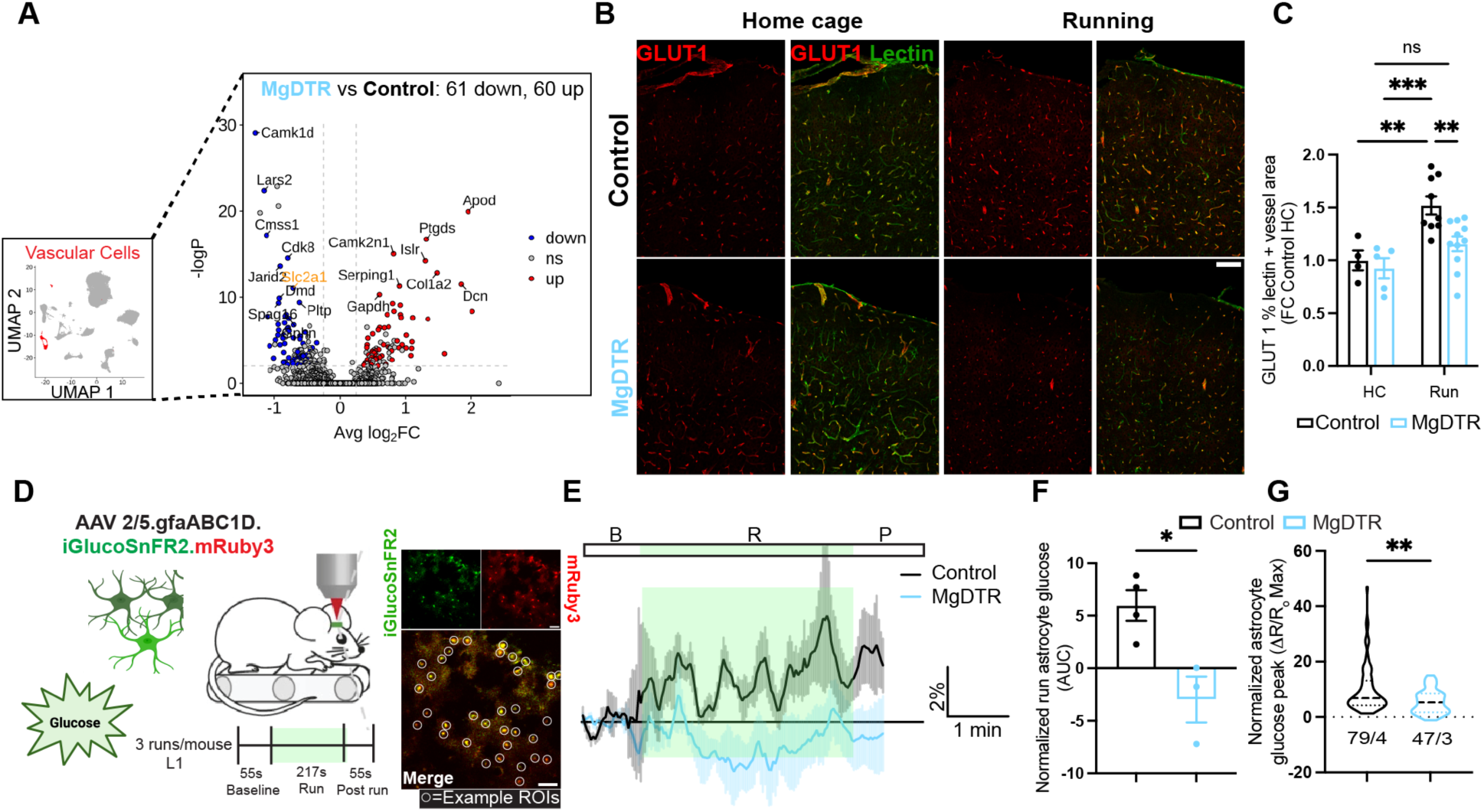
Microglia actively regulate brain glucose entry. (**A**) Volcano plot of the Sn-RNA seq vascular cell cluster (red in UMAP, left) with genes upregulated (red), non-significant (grey), or downregulated (blue) in MgDTR mice including *Slc2a1* (*Glut1*, orange). (**B**) Representative images of GLUT1 (red) staining in motor cortical lectin+ blood vessels of in control or MgDTR mice in HC or Running conditions quantified (**C**) as % lectin area covered in GLUT1 per mouse, normalized to Control HC mean (N_CHC_=4 mice, N_MHC_=5 mice, N_CRun_=9 mice, N_MRun_=11 mice). (**D**) Schematic of running paradigm for iGlucoSnFR2 imaging in L1 astrocytes and (right) representative image of astrocytic sensor expression. (**E**) Mean animal 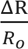 L1 astrocytic intracellular glucose traces during baseline (B), run (R), and post run (P) phases in control (black) and MgDTR (blue) mice (N_Cntr_=4 mice, N_MgDTR_=3 mice). Shading represents +/- S.E.M. (**F**) Run AUC per mouse and (**G**) peak iGlucoSnFR2 signal per astrocyte (N_Cntr_=79 astrocytes from 4 mice, N_MgDTR_=47 astrocytes from 3 mice). Unpaired two-tailed student’s *t* test (F) and (G). Two-way ANOVA with Tukey’s post hoc test for (C). *P<.05, **P<.01, ***P<.001; ns= not significant. Scale bar: 50 µm for all images. Illustration in (D) created with BioRender.com.

The purported first step in the ANLS is activity-dependent uptake of glucose from blood into astrocytic endfeet, which travels through capillary and astrocytic GLUT1 ^1,8^. To test whether deficits in endothelial GLUT1 were associated with reduced uptake of glucose into the ANLS, we expressed the intracellular fluorescent glucose sensor iGlucoSNFR2 ^65^ in motor cortical astrocytes in MgDTR mice and controls using AAV 2/5.GFAP.(cyto).iGlucoSNfR2.mRuby3 and compared astrocytic glucose fluxes during running (**Figure 6D**). In mice with microglia, running led to an increase in intracellular glucose in astrocytes, mimicking the effects of glutamate *in vitro* ^39^ and blood glucose tracing during whisker stimulation *in vivo* ^7,9^. However, in MgDTR mice, integrated astrocytic glucose failed to increase above baseline during motor training and there was an overall decrease in peak relative sensor fluorescence. These findings are consistent with deficits in endothelial GLUT1 in the absence of microglia (**Figures 6E-6G**).

Brain macrophage derived vascular endothelial growth factor A (VEGFa) is known to regulate brain endothelial GLUT1 expression and glucose uptake in pathological conditions ^66^. Although one of the most enriched GSEA pathways in motor-trained microglia from our TRAP-seq dataset was “positive regulation of vascular endothelial growth factor production,” VEGFa itself was not significantly upregulated (**Figure S4F**). However, the mRNA encoding growth factor *Cyr61* (*Ccn1*), a known VEGFa regulator ^67^, was significantly increased in microglia from trained mice (Log_2_FC= 1.3326, P_adjusted_= .0021) (**Data S5**). Expression of *Cyr61* directly correlates with levels of *Glut1* ^68^ and conditional knockout of *Cyr61* in intestinal stem cells leads to reduced enterocyte GLUT1 and blood glucose absorption ^69^. Furthermore, *Cyr61* is an immediate early gene, and thus has the capacity for rapid synthesis in response to a stimulus ^70^. To confirm that CYR61 is increased in microglia at the protein level during motor activity, and to exclude an effect of other CX3CR1 expressing resident border-associated macrophages (BAMs) ^71^, we performed flow cytometry on enriched motor cortical cells from rotarod running and HC mice, staining for CYR61 along with cell-type specific markers (**Figures 7A and S5A**). To our surprise, we found reduced CYR61 protein levels in motor-trained microglia (CD64^+^/CD206^-^/MHC2^-^) (**Figures 7B and 7C**). In contrast, CYR61 was unchanged in two BAM populations (CD64^+^/CD206^+^/MHC2^-^ or CD64^+^/CD206^mid^/MHC2^High^) ^72^ (**Figures S5B-S5E**). Because CYR61 is a secreted protein ^70^, we asked whether the increase in ribosome-bound *Cyr61* mRNA we detected might reflect new synthesis used to restore internal CYR61 protein stores that are rapidly secreted and depleted on demand. CYR61 is known to be sensitive to brefeldin A, a fungal metabolite that blocks protein secretion ^73^. Therefore, to determine whether CYR61 is actively secreted by microglia during motor activity *in vivo*, we injected mice with brefeldin A, which has been to shown to block cytokine secretion *in vivo* ^74,75^, 2.5 hours prior to the rotarod task, and conducted flow cytometry as before (**Figure 7D**). Here, we detected an increase in microglial CYR61 protein in running compared to HC mice, with no significant change in BAM CYR61 (**Figures 7E, 7F and S5F-S5I**). Taken together, these findings suggest that microglia CYR61 is both synthesized and secreted in response to motor activity.

**Figure 7.**
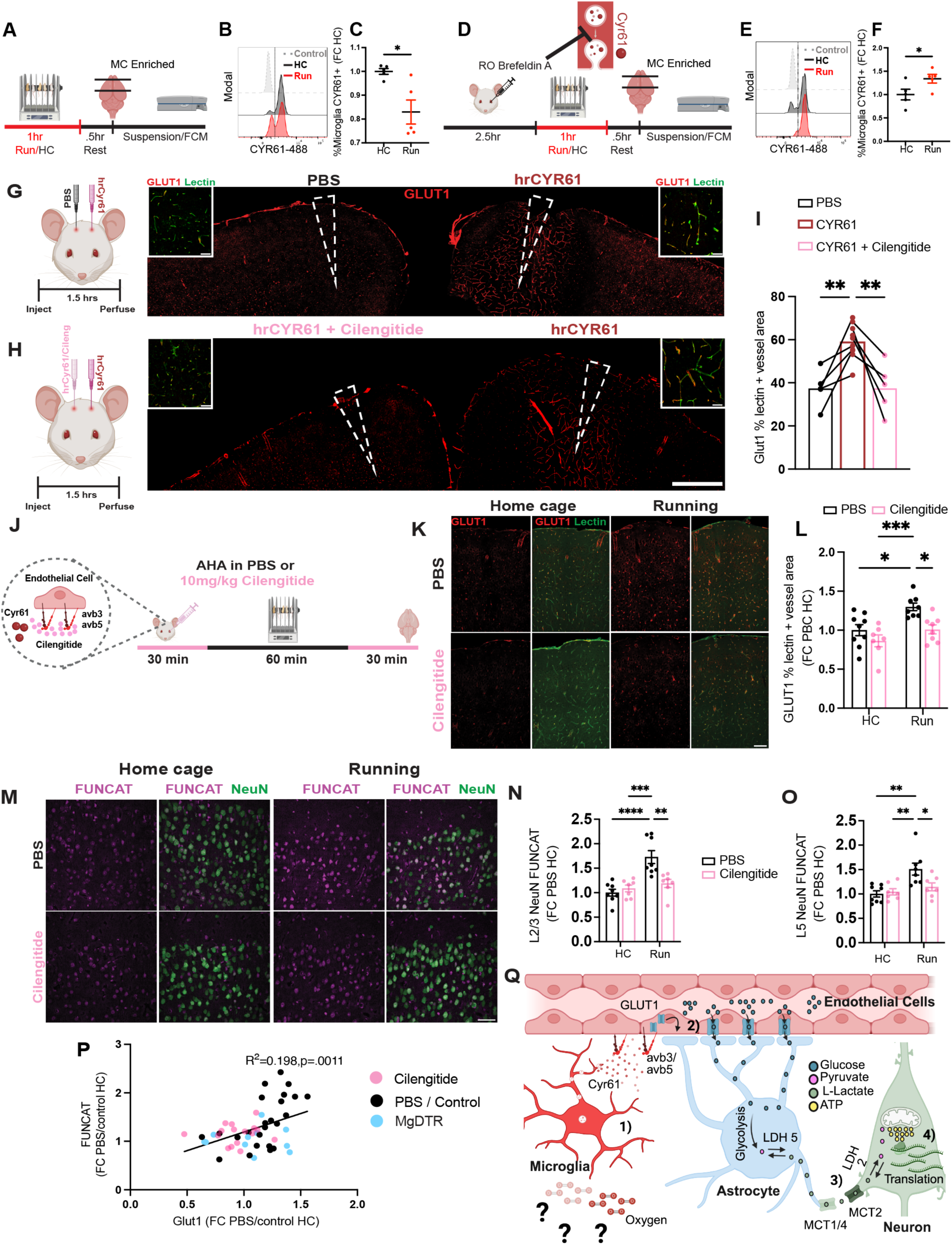
Cyr61-integrin signaling regulates endothelial GLUT1 and *de novo* translation in neurons. (**A**) Schematic of rotarod flow cytometry protocol. (**B)** Representative histogram of gated microglia (Figure S5A) CYR61 staining from HC (black), Running (red), and fluorescence minus one (FMO, grey) control group. (**C**) FMO subtracted Cyr61 % positive cells per mouse normalized to HC mean (N=5 mice for both groups). (**D** to **F**) Similar to (A) to (C) but for an experimental setup in which mice were injected with brefeldin A (which block vesicle secretion as illustrated, top) 4 hours prior to perfusion. (**G**) (Left) Schematic of experiment in which the left and right motor cortices of mice were injected with PBS and .8μg hrCYR61 respectively and perfused 1.5 hours later. (Right) representative image showing GLUT1 (red) immunostaining in both cortices and approximate injection location (white triangle). (**H**) Similar to (G) but for experiments in which the left cortex was injected with .8μg hrCYR61 and 1μg cilengitide. (**I**) Quantification of GLUT1 % area in lectin+ blood vessels in motor cortex after PBS (black), Cyr61 (brown), or Cyr61+Cilengitide (pink) (N=5 mice/experiment). (**J**) Schematic of *in vivo* FUNCAT experiment in which mice were injected with AHA in pbs or 10mg/Kg cilengitide 30 mins before motor training which (left illustration) blocks the receptors of Cyr61, avb3 and avb5. (**K**) Immunostaining of motor cortical GLUT1 (red) in Lectin+ blood vessels (green) and (**L**) quantification of percent of blood vessel area covered in GLUT1, normalized to PBS HC mean animal value (N_PHC_=9 mice, N_CHC_=7 mice, N_PRun_=8 mice, N_CRun_=7 mice). (**M**) Representative images of L2/3 MC neuronal (green) FUNCAT (purple) labeling and quantification in (**N**) L2/3 and (**O**) L5 per animal normalized to PBS HC mean animal value (N_PHC_=8 mice, N_CHC_=7 mice, N_PRun_=8 mice, N_CRun_=7 mice). (**P**) Correlation between increased FUNCAT labeling and blood vessel GLUT1 staining for all experiments in which mice were labeled for FUNCAT and GLUT1, normalized to within experiment HC control group (either PBS injected Wild-type or undepleted Control mice). Colored dots represent the genotype or experimental condition replicates came from (N= 51 mice). (**Q**) Proposed model for microglia metabolic coupling. Unpaired two-tailed student’s t-test for (C) and (F). Repeated measures mixed effects model with Sidak’s post hoc test for (I); Two-way ANOVA with Tukey’s post hoc test for (L), (N), and (O); Simple linear regression for (J). Drug x condition interactions for (N) and (O) respectively F(1,26)=10.88, 5.905, **P=.0028,*P=.023. *P<.05, **P<.01, ***P<.001 ****P<.0001. Scale bar: 200 µm for (H) and 50 µm for (K) and (M). Illustrations (G), (H), (J), and (Q) created with BioRender.com.

Although our results establish that running induces changes to ribosome bound *Cyr61* mRNA and protein in microglia, the secretion of CYR61 in an activity-dependent manner may be a generalized response undertaken by multiple vascular-associated cell types. To determine the specificity of microglia CYR61 secretion in our paradigm, we repeated our CYR61 flow cytometry assay as before, evaluating whether *Cyr61* expressing astrocytes or endothelial cells^76^ actively secrete CYR61 during running (**Figure S5J**). We detected intracellular CYR61 in both astrocytes and endothelial cells, but we observed no difference in CYR61 levels in either cell type between running and HC mice (**Figures S5K-S5M**). Taken together, these results suggest that CYR61 secretion is likely a microglia-specific response to increased activity.

Next, to test whether CYR61 can directly regulate brain endothelial GLUT1 expression, we injected recombinant human CYR61 (rhCYR61) into the right motor cortex of wild-type mice and compared endothelial GLUT1 expression with the contralateral saline-injected left motor cortex (**Figure 7G**). 1.5 hours post injection, levels of vascular GLUT1 were significantly increased in rhCYR61-injected cortices, particularly around the injection site (**Figures 7G and 7I**). Co-injection of rhCYR61 with cilengitide, a potent inhibitor of the endothelial CYR61 integrin receptors avb3/avb5 ^70,77^ prevented the increase in endothelial GLUT1 (**Figures 7H and I**). Thus, CYR61-integrin signaling can rapidly facilitate increases in brain endothelial GLUT1 expression.

Finally, we tested whether CYR61-integrin signaling is required for activity-dependent GLUT1 expression in endothelial cells and ultimately, *de novo* protein synthesis in neurons. We conducted *in vivo* FUNCAT labeling while blocking CYR61-integrin signaling by co-injecting cilengitide (10 mg/Kg) with AHA, 30 mins prior to the rotarod task (**Figure 7J**). Similar to results in MgDTR mice, injection of cilengitide blocked the increase in endothelial GLUT1 expression following rotarod running (**Figures 7K and 7L**). In addition, cilengitide injection blocked the motor task-induced increases in neuronal *de novo* protein synthesis in both L2/3 and L5 (**Figures 7M-7O**). Moreover, we observed a direct correlation between increases in endothelial GLUT1 levels and neuronal *de novo* protein synthesis (**Figure 7P**). Synthesizing all our results, we propose a model in which microglia regulate local brain metabolism in an activity-dependent manner through the following mechanism. First, increased activity causes microglia to rapidly secrete CYR61, that is eventually restored through *de novo* protein synthesis. Second, acting as a growth factor, CYR61 binds to endothelial integrin receptors initiating an increase in GLUT1. Third, increased GLUT1 allows for on-demand glucose entry into the ANLS elevating neuronal ATP levels within 30 mins. Fourth, and finally, the increased supply of ATP is utilized for the metabolically costly process of protein synthesis in neurons, which is elevated within 90 mins (**Figure 7Q**).

## Discussion

Overall, our findings reveal a novel mechanism involving the coordination between multiple brain cell types, allowing for the maintenance of tissue homeostasis in the face of sustained changes in metabolic demand. We propose that microglia are at the helm of this homeostatic apparatus and synchronize the system at its primary level: glucose entry into the brain. Our results support and provide a rationale for recent PET imaging studies that suggest that adult brain metabolic connectivity and glucose entry are reduced in either the absence of microglia or after microglia-specific mutations ^78–82^. Furthermore, our findings argue against a cell-autonomous model of microglia involvement in brain glucose uptake ^82^ but rather for a non-cell autonomous model in which microglia regulate astrocyte glucose uptake from blood, in agreement with Zimmer and colleagues ^83^.

Although our findings suggest that microglia control of brain metabolism is primarily at the level of endothelial cells, it does not exclude other lines of communication with other cells. Indeed, adenosine has recently emerged as an important regulator of astrocyte glycolysis and lactate shuttling ^40^ and microglia are thought to produce adenosine in an activity-dependent manner to dampen (or perhaps feed) hyperactive circuits ^21^. In addition, in the absence of microglia, oligodendrocyte clusters arise with dysregulated lipid metabolism, which is tied to deficits in microglial TGF-β ^84^. Thus, in future studies it will be important to explore and compare different brain states and loci at which microglia regulate brain metabolism.

The trigger for activity-dependent CYR61 release from microglia is presently unknown. *Cyr61* is a canonical hypoxia responsive gene and cultured brain macrophage *Cyr61* mRNA increases in response to hypoxia ^60^. However, accounting for the rapid nature of the observed effects, our model proposes *de novo* microglial CYR61 mRNA translation is used to restore the pool of CYR61 that was secreted. Regarding secretion, changes in oxygen have been shown to regulate CYR61-CAV1 interactions, caveolae formation, and CYR61 secretion^73^. Running is known to induce a transient neuroprotective brain state termed “functional hypoxia” in the hippocampus and cortex ^85,86^. Thus, local changes in oxygen in the motor cortex elicited by motor activity may trigger CYR61 release. However, microglia express a variety of receptors that are used to sense neuronal activity and neuromodulation ^87^. Therefore, a more direct mechanism whereby microglia sense and respond to byproducts of neuronal activity should also be explored.

Similarly, the exact mechanism(s) by which CYR61-integrin receptor signaling leads to rapid increases in endothelial GLUT1 levels and/or membrane trafficking remain to be determined. There are at least three potential mechanisms for how integrin receptor activation leads to increased endothelial GLUT1. One possibility is that integrin receptor activation could induce the transcription of GLUT1. Another possibility is that CYR61-integrin receptor signaling could influence the translation of GLUT1 independent of transcription. Finally, integrin receptor activation could regulate the degradation/recycling of GLUT1. CYR61-integrin binding is known to activate transcriptional activator YAP ^69,88^, which has been shown to increase the expression *Slc2a1* (Glut1)^68^. However, considering the temporal confines of our model, mechanisms operating at either the translational or recycling level(s) appear to be more likely. For example, CYR61-integrin signaling is known to activate Akt^89,90^, which has been shown to promote GLUT1 trafficking and glucose transport by other growth factors^91^. With respect to the temporal features of our model, it should be noted that prior to the forced running paradigms, the experimental mice run freely in their cage. Therefore, subtle deficits in endothelial GLUT1 may be already apparent in MgDTR mice at baseline, which are fully evoked when metabolic demand increases during forced running. This possibility is supported by a significant main effect of genotype on endothelial GLUT1 levels. Although blocking the receptor of CYR61 immediately prior to running is sufficient to prevent running-induced increases in endothelial GLUT1 and neuronal protein synthesis, further work is needed to fully disentangle the precise temporal features of microglia metabolic coupling.

It is becoming increasingly clear that a number of neurodegenerative diseases involve both hypometabolic states ^1^ and proteostasis failure ^92^. Considering the large metabolic demand required for mRNA translation, a direct connection is compelling. Furthermore, in Alzheimer’s disease, a decrease in capillary GLUT1 is well established ^25^, as are prominent phenotypic changes in microglia ^93^. It is tempting to speculate that microglial transcriptional changes, induced by disease states, limit their capacity for metabolic coupling, which in turn limits neuronal protein synthesis and, ultimately, memory.

### Limitations of the Study

There are several limitations of this study to be addressed. First, although technical advancements in protein labeling using techniques such as in vivo FUNCAT^37^ now allow for the visualization of activity-dependent protein synthesis within 2 hours of a stimulus, it is still unclear exactly when activity converts to new protein synthesis *in vivo*, when new ATP is essential, and when over the course of 1.5 hours microglia are required for this process. New sensors providing real-time readouts of *de novo* protein synthesis that are amenable to *in vivo* imaging would allow for a more precise understanding of temporal features of the proposed model. Next, although it is known that motor cortical protein synthesis^35^ and microglia^16^ are independently important for motor learning, it is still unclear whether microglia metabolic coupling is necessary for both activity-dependent protein synthesis and learning or just the former. Finally, our findings emphasize the importance of endothelial GLUT1 regulation to meet the metabolic demands for new protein synthesis, but we provide only one mechanism for its regulation in an activity-dependent manner. It is conceivable that endothelial GLUT1 is regulated through multiple mechanisms in numerous contexts and in different brain regions. Likewise, it remains to be seen whether microglia metabolic coupling is a feature of motor cortical activity, or whether it functions in other regions relevant for other types of behavior.

## Acknowledgments

We would like to thank Dr. Stephen Zhang for his insightful comments on the manuscript; Maggie Donohue for her expertise in animal husbandry, colony management, and protocol organization; the NYU microscopy core and Michael Cammer in particular for their expertise in microscopy and image analysis; Dr. Juan Lafaille and for his continual support and advice; and the NYU cytometry and cell sorting core facility for help with flow cytometry. Illustrations (specified in figures) were created with BioRender.com.

## Author contributions

Conceptualization: DA, AMA, MMO, HTE, WBG, EK

Methodology: DA, AMA, HE, MMO, MZ, RS, MCS, OM, WL, DY, AXG

Investigation: DA, AMA, EC, HTE, MMO, MZ, RS, MCS, OM, WL, NDS

Visualization: DA, AMA, MMO, MZ, RS

Funding acquisition: DA, MVC, RCF, WBG, EK, SAL

Supervision: DA, WBG, EK

Writing – original draft: DA

Writing – review & editing: AMA, HTE, MMO, MC, MCS, RCF, WBG, EK, SAL

## Data and materials availability

Omic data will be deposited on the gene expression omnibus (GEO) by time of publication.

---

**Data S1. (separate file)**

List of Hallmark gene sets discovered (FDR q<.01) for clusters from Sn-RNA seq (see Fig. 1C) via GSEA.

**Data S2. (separate file)**

List of gene ontology biological pathways (GOBP) gene sets discovered (FDR q<.01) for clusters from Sn-RNA seq (see Fig. 1C) via GSEA.

**Data S3. (separate file)**

List of DEGs from microglia Sn-RNA seq experiment (see Fig. 1C) for large cell type groups (see methods section, “*Differential gene expression and pathway analysis”*).

**Data S4. (separate file)**

List of DEGs from microglia Sn-RNA seq experiment (see Fig. 1C) for all clusters.

**Data S5. (separate file)**

List of DEGs from microglia TRAP-seq experiment (see Fig. 5K). Tab 1 is all genes P<.05. Tab 2 is all genes P_adjusted_<.05. For all analyses only DEGs with P_adjusted_<.05 were considered significant.

## Materials and Methods

### Animals

All animal protocols were reviewed and approved by the New York University Animal Care and Use Committee or by the Institutional Animal Care and Use Committee of Shenzhen Graduate School, Peking University (approved protocol 11514), and adhered to institutional guidelines. Mice were provided with food and water ad libitum and were maintained in a 12 h– 12 h light–dark cycle at New York University at a stable temperature (78 °F) and humidity (40– 50%). Cx3cr1^CreERT2^ (JAX:020940 for all experiments except snRNA-seq, which used 021160), ROSA26iDTR (iDTR; JAX:007900), tdTomato (Ai14; JAX:007908), Camk2a^Cre^ (T29-1; JAX:005359), and floxed TRAP (EGFP-L10a; JAX:024750) mice were purchased from Jackson labs. Thy1.2 GCaMP6s line 3^95^ were provided by Dr. Wen-Biao Gan. Cx3cr1^CreERT2^ and iDTR mice were crossed to generate heterozygous MgDTR mice (Cx3cr1^CreERT2/+^:R26^DTR/+^) and were compared to Cx3cr1^CreERT2/+^:R26^+/+^ littermates. Cx3cr1^CreERT2^ and EGFP-L10a mice were crossed to generate heterozygous microglia TRAP (Cx3cr1^CreERT2/+^:R26^EGFP-L10a/+^). Camk2a^Cre/+^ mice were crossed with Ai14 mice to generate heterozygous microglia tdTomato mice (Camk2a^Cre/+^:R26^tdTomato/+^). Male and female mice aged 2-6 months were used in all experiments, and all litters were randomized into experimental groups.

### Drugs, chemicals, and recombinant proteins

Tamoxifen (Sigma T5648) was dissolved in corn oil (Sigma) at 20mg/ml. Mice at least 4 weeks old were gavaged with .5mg/g tamoxifen oil 2x two days apart. Diphtheria toxin (DT; Sigma, D0564) was reconstituted to 1mg/ml in sterile PBS and stored at −80^0^C. For microglia depletion, DT was further diluted to .01mg/ml, immediately prior to use, and 100µl were injected per mouse intraperitoneally (IP) 1x/day for three consecutive days^16^. DT was injected at least 3.5 weeks after tamoxifen gavage to allow for repopulation of unrecombined peripheral cells while preserving DTR expression in long-lived microglia^16^. Oligomycin A (Sigma, 75351) was reconstituted in Dimethyl sulfoxide (DMSO) and then further diluted to 40uM in artificial cerebral spinal fluid (ACSF). To block ATP synthesis *in vivo*, oligomycin A was applied to an open cranial window allowing penetration to the underlying cortical region. AR-C155858 (Tocris 4960) was reconstituted in DMSO and then further diluted to 50µM in ACSF with 50mM L-lactate (Sigma, L7022). The AR-C155858 + L-lactate solution was added to an open cranial window as with oligomycin A. L-azidohomoalanine (AHA; Vector Labs CCT-1066) was diluted to 50mg/ml in sterile PBS and stored at −20^0^C. Prior to use, AHA was further diluted to 25mg/ml and injected retro-orbitally (50mg/kg). α-cyano-4-hydoxcinnamic acid (4-CIN; Sigma, C2020) was diluted in PBS (for *in vivo* FUNCAT experiments) or ACSF (for *in vivo* imaging) to 12mM, brought to a pH of 8 to fully dissolve, and returned to a pH of 7.4. For FUNCAT experiments, mice were IP injected with 4-CIN at 90mg/kg 15 minutes (mins) prior to motor training. For *in vivo* imaging experiments, 1.2mM 4-CIN was applied directly to the cortex under a cranial window. For lactate rescue *in vivo* FUNCAT experiments in MgDTR mice, L-lactate diluted in PBS was prepared fresh for each experiment and mice were injected (1g/kg) IP immediately prior to and after training. Recombinant human CYR61 Fc chimera protein (R&D, 4055-CR-050) was diluted in PBS and 800ng was stereotactically injected into motor cortex (see below). Cilengitide trifluoroacetic acid salt (cilengitide; Sigma, SML1594) was diluted in PBS, combined with AHA, and injected (10mg/kg) retro-orbitally for *in vivo* FUNCAT. For stereotactic co-injections with hrCYR6, 1μg cilengitide was used.

### AAV injection

Newborn mice (P1-4) received 400nl intraparenchymal injections of either AAV-2/1.hSyn1.(cyto).iATPSnFR2.A95A.A119L.mCherry (P1-3, 1×10^13 GC/mL)^27^, AAV-2/5.gfaABC1D-eLACCO2.1 (P3-4, 8.9×10^12 GC/mL)^31^, or AAV-2/5.GFAP-iGlucoSnFR2.mRuby3 (1×10^13 GC/m)^65^ (P3-4) through the skull above the motor cortex using a 5ul syringe fitted with a 33 gauge needle (Hamilton, 65460-03) after cryoanesthesia. The approximate motor cortex was identified as the region just rostral of the lateral ventricles, which could be identified through the neonatal translucent scalp. The needle was inserted until the full bevel length was within the skull (approximately .987mm). We noticed astrocytic expression improved when AAVs were injected later in development (P3-P4 vs P1-2). The virus was injected at a rate of 16.6 nl/s using robot stereotaxic microinjection pump (Neurostar), after which the needle was kept in the parenchyma for an extra 10s to reduce backflow. AAV-2/1.hSyn1.(cyto).iATPSnFR2.A95A.A119L.mCherry and AAV-2/5.GFAP-iGlucoSnFR2 were supplied by Janelia Constructs. AAV-2/5.gfaABC1D-eLACCO2.1 was generously provided by Dr. Robert Campbell.

### hrCYR61 Injection

Mice were anesthetized with isoflurane. A midline incision was made over the skull, and two holes were drilled over the left and right motor cortices using an automatic drill (NeuroStar) based off the measured head tilt and injection coordinates. The right motor cortex was stereotactically injected with 800nl hrCYR61 and left motor cortex with equal volumes of either PBS or hrCYR61 + cilengitide using the following coordinates: AP 1.25, ML +/- 1.0, DV 1.6. The needle was inserted at a rate of 1mm/min and was inserted until .2mm below the intended injection site. The needle was left beneath the injection site 3 mins prior to injection after which it was moved to the correct coordinates. After injection, the needle remained in the injection site for an additional 5 mins to minimize backflow. The needle was then removed at 1mm/min, and the contralateral hemisphere was then injected. The order of hemisphere injection was alternated for each mouse. After the second injection, the scalp was glued together and the mouse recovered on a heating pad. Mice were perfused 1.5 hours after the second injection.

### Surgical preparation for imaging awake, head-restrained mice

Sensor imaging was carried out in awake, head-restrained mice through an acute open skull window. MgDTR mice and controls were prepared 1-2 days after the 3^rd^ injection of DT. Surgery preparation for awake animal imaging includes attaching the head holder and creating open cranial window over the primary motor cortex (∼0.5 mm anterior from bregma and ∼1.2 mm lateral from the midline). Specifically, mice were deeply anaesthetized with an intraperitoneal injection of ketamine (100 μg g^−1^) and xylazine (10 μg g^−1^). The mouse head was shaved, and the skull surface was exposed with a midline scalp incision. The periosteum tissue over the skull surface was removed without damaging the temporal and occipital muscles. Mice were then screened using a fluorescent microscope for adequate expression of the sensor through the skull and a mark was made over the primary motor cortex, which corresponded to the maximal sensor expression. Mice with clear expression of sensor were implanted with head bars.

A head holder, composed of two parallel metal screws, was attached to the mouse’s skull to help restrain the mouse’s head and reduce motion-induced artefact during imaging. A thin layer of cyanoacrylate-based glue was first applied to the top of the entire skull surface, and the head holder was then mounted on top of the skull with dental acrylic cement (Lang Dental Manufacturing Co.) such that the marked skull region was exposed between the two bars. Precautions were taken not to cover the marked region with dental acrylic cement.

After the dental cement was completely dry, the head holder was screwed to two metal cubes that were attached to a solid metal base, and a cranial window was created over the previously marked region. 1.8mm circular region was outlined on the scull by gently drilling with a 1.8mm trephine (224RF-023-HP, Meisinger) inserted in a high-speed drill. The trephine was then replaced with a circular drill bit to carefully shave the 1.8mm region until the bone could be removed. Once removed the bone was replaced with a custom made 1.8mm glass coverslip (0 thickness, Potomac Laser). ACSF was periodically applied to the skull during drilling to minimize damage caused by heat from the drill. For preps not requiring application of metabolites or drugs, the cover slip was glued completely. For preparations in which drugs/metabolites were applied to the cranial window, a custom cut square cover slip was alternatively used as a cranial window and only 3 sides of the window were glued, keeping one side open and covered in ACSF until application of the drug/metabolite. Imaging experiments were started within 24 h after window implantation. Awake mice were head restrained in the imaging apparatus, which sits on top of a custom-built free-floating treadmill.

### Treadmill training and *in vivo* two-photon microscopy

A custom-built free-floating treadmill (96 × 56 × 61 cm dimensions) was used for motor training. This free-floating treadmill allows head-fixed mice to move their forelimbs freely to perform motor running tasks. Mice were positioned on a custom-made head holder device that allowed the micro-metal bars (attached to the mouse’s skull) to be mounted and for the base of the device to be positioned below the belt in contact with the microscope stage. During motor training, the treadmill motor (Dayton, Model 2L010) was driven by a DC power supply (Extech). At the onset of a trial, the motor was turned on and the belt speed gradually increased from 0 cm s^−1^ to 8 cm s^−1^ within ∼3 s.

For MgDTR iATPSnFR2 imaging, mice were run 2-3x for 100 frames (“short runs”, 2.173 sec/frame, 2µS pixel dwell time) with a 1 min inter-run rest followed by 1x 250 frame “long run” in L2/3 (150-250µM below the pial surface) motor cortex. For both short and long runs, a 25-frame baseline was taken immediately prior to commencing running. For short runs, a 25-frame “post run” period was imaged (150 frames total). For long runs, a 150-frame “post run” period was imaged (425 frames total). After the end of the long run’s post run period, mice were rested for 5 mins before repeating the protocol in L1 (25-100µM below the pial surface). In experiments assessing the effect of 4-CIN on ATP, L1 neuropil of wild-type mice was first imaged through an open cranial window covered in ACSF during short and long runs. Then 1.2mM 4CIN was applied to the open window and was incubated for 30 min before restarting the protocol. For iGlucoSnFR2 imaging, 2-3 150-frame short runs were conducted per mouse in L1.

iATPSnFR2 in neurons (hSyn1 promoter) and iGlucoSnFR2 in astrocytes (GFAP promoter) were imaged with an Olympus FVMPE-RS multiphoton laser scanning microscope (galvo scanner mode) equipped with two tunable lasers (Insight x3-OL and MaiTai HP DS-OL) with a beamsplitter set to 690/1050. All experiments were performed using an Olympus 25x N.A.

1.05 XLPLN25XWMP2 objective immersed in either ACSF or ultrasound gel (Aquasonic Clear). For iATPSnFR2 experiments, iATPsnFR2 was excited at 930nm and mCherry was excited at 1075nm. For iGlucoSnFR2 experiments, iGlucoSnFR2 was excited at 930nm and mRuby3 was excited at 1080nm. Green (iATPSnFR2 or iGlucoSnFR2) emission was collected through a 538/40 filter and red (mCherry/mRuby3) emission was collected through 610/35 into GaAsP detectors. For experiments with assessing the effect of Oligomycin A on iATPSnFR2 in neurons, wild-type mice were anesthetized with ketamine/xylazine (KX) and an open cranial window was covered with ACSF. While under the microscope, ACSF was exchanged with 40 μM Oligomycin A and L1 neuropil ATP was imaged for 425 frames. All images were acquired using sequential line scanning with a 512 x 512 pixel aspect ratio at 2µsec/line, 2.12msec/line, and 2.173sec/frame.

eLACCO2.1 around astrocytes (gfaABC1D promoter) in layer 1 (25-100µM below the pial surface) were imaged with an Olympus Fluoview 1000 two-photon system (tuned to 930 nm) equipped with a Ti:Sapphire laser (MaiTai DeepSee, Spectra Physics) 50 frames before, 150 frames during each run, and for 50 frames after each run, after which scanning was terminated for 60s until the next baseline was taken. For experiments assessing the effect of AR-C155858 and L-Lactate on eLACCO2.1 signal around astrocytes, WT mice were anesthetized with KX and an open cranial window was covered with ACSF. While under the microscope, ACSF was exchanged with 50mM L-Lactate + 50µM AR-C155858 and L1 extracellular lactate around astrocytes was imaged for 500 frames. All experiments were performed using a x25 objective (numerical aperture 1.1) immersed in an ACSF solution and with a x1.7 digital zoom. All images were acquired at 1.109 frames/second (2µsec pixel dwell time) with a 512 x 512 pixel aspect ratio.

All experiments except sensor validation in Fig. S2H were conducted in unanesthetized mice. The dura was left intact for all experiments.

### Rotarod *in vivo* FUNCAT

*In vivo* FUNCAT was conducted as described^37^ in combination with rotarod training with minor adjustments. Mice were injected with AHA retro-orbitally (50mg/kg) and put back in their home cage for 30 mins to allow AHA incorporation into the brain. After a 30-min rest, “running” mice were put on a rotarod (476000, Ugo Basile) set to a fixed speed of 12RPM for 1 hour, while “home cage” (HC) mice were left in their cage. Mice that fell off the rod were immediately placed back on the rod. After 1 hour of training, mice were returned to their home cage for 30 mins after which they were perfused with PBS, followed by 4% paraformaldehyde (PFA). After an overnight post fixation step at 4°C, 40µM coronal brain slices were generated from regions containing the motor cortex (Bregma 1.94-1.0). Sections were added to 24 well plates (maximum 3 slices/well) and permeabilized/blocked with 5% bovine serum albumin (BSA), 5% normal goat serum (NGS, 005-000-121 Jackson Immuno), and .3% Triton x100 in PBS for 2 hours at room temperature (RT) on an orbital shaker. After block/permeabilization sections were washed 3x 10 mins with PBS. Next copper-catalyzed click reactions were conducted using 500µL per well of components from either of the following kits (C10269, ThermoFisher or CCT-1263, Vector Labs) and Alexa Fluor 647 Alkyne, Triethylammonium Salt (A10278 ThermoFisher) or AZDye 647 Alkyne (CCT-1301, Vector Labs) (diluted to 1mM in DMSO) added in the following order per well: 1) 438.75µL cell reaction buffer 1x 2) 50µL additive 3) 1.25µL Alkyne 647 4) 10µL CuS04 solution. Components were vortexed after each component addition. Reactions proceeded overnight at 4°C on an orbital shaker. The next day sections were washed 5x 5min with .5mM ethylenediaminetetraacetic acid (EDTA) and 1% Tween-20 in PBS on an orbital shaker. Sections were then mounted with Fluoromount-G with DAPI (00-4959-52, ThermoFisher) or stained with cell type specific markers using immunofluorescence. All mouse FUNCAT signal was compared to no AHA controls injected with PBS to ensure FUNCAT signal was above background. Slices with cuts in the cortex were avoided because of increased non-specific click labeling.

### Immunofluorescence/Imaging

If not previously blocked/permeabilized with prior to FUNCAT, free floating sections were blocked/permeabilized with 5% normal goat serum (NGS, 005-000-121 Jackson Immuno), and .3% Tritonx100 in PBS for 2 hours at RT on an orbital shaker and washed 3x 10 mins in PBS. Sections were then stained overnight at 4°C in the following primary antibodies or dyes diluted in PBS with .1% Tritonx100 (PBST): guinea pig anti-NeuN (1:1000, 266 004, Synaptic Systems), chicken anti-IBA1 (1:500, 234 009, Synaptic Systems), rabbit anti-IBA1 (1:1000, 197-19741, Wako), Rabbit anti-GLUT1 (1:300, ab115730, abcam), and FITC Tomato Lectin, (1:300, FL-1171-1, Vector Labs). The following day sections were washed 3x 10 mins in PBST and stained in the following secondary antibodies all at 1:500: goat anti-guinea pig Alexa Fluor 488 (A-11073, ThermoFisher), goat anti-chicken Alexa Fluor 568 (A-11041, ThermoFisher), or goat anti-rabbit Alexa Fluor 568 (A-11011, ThermoFisher) for 1.5 hours at RT. Samples were then washed 2x 10 mins with PBST and 1x with PBS before mounting as with FUNCAT.

Images were acquired using a Zeiss LSM 800 confocal microscope equipped with a 20x objective for FUNCAT % area cation, GLUT1 % area quantification, or microglia morphological quantification. A 40x/1.3 N.A. oil emersion objective was used for neuronal or microglial FUNCAT quantification. Tiles were stitched using Zen Blue software (Zeiss). Identical microscope settings (e.g. imaging depth, z-step size, laser power, gain, optical zoom, offset, filters) were used for images in which fluorescent intensity, % area, or cell morphology were quantified and compared. 40x images from layer 2/3 were taken approximately 150-400µM beneath the pial surface (measured using Zen Blue software) and 550-700µM beneath the pial surface for L5. Regions were chosen immediately above the dorsal peak of the corpus callosum anterior forceps.

### Microscopy image analysis

#### Metabolic sensors

Time course images were imported to FIJI (ImageJ) for analysis. To correct for x,y plane movement during running, image channels were duplicated, merged (for dual color sensors), and registered to the first frame of the image using the plugin TurboReg^96^. For iATPSnFR2 imaging in L1 neuropil, a single large region of intensity (ROI) covering the brightest regions of tissue (avoiding dark blood vessels) was generated for each run. For L2/3 Somatic iATPSnFR, L1 dendritic GCaMP6s, L1 astrocytic eLACCO2.1, and L1 astrocytic iGlucoSNFR2, individual ROIs were manually drawn around somas, dendrites, or astrocytes (including processes) and mean fluorescence intensity (MFI) was calculated for each ROI. For dual-fluorophore sensors (iATPSnFR2-mCherry and iGlucoSnFR2-mRuby3), green/red MFI ratios (R) were calculated at each timepoint and filtered by taking a 5-frame moving average, as has previously been done for metabolic sensor imaging,^97^ to generate 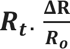 was generated by calculating the % change of ***R***_***t***_ from a 25-frame pre-run baseline (***R***_***o***_) for each individual ROI:

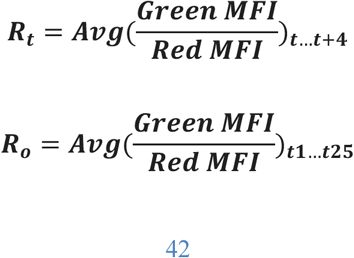

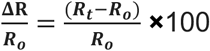

For single fluorescent sensor imaging (GCaMP6s or eLACCO2.1), 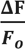 was calculated as with 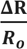 without taking a ratio of fluorophores. Additionally, for GCaMP6s ***F***_***o***_ represented the average MFI of the first 10 frames and for eLACCO2.1 ***F***_***o***_ represented the average MFI of the first 50 frames. A moving average was not used for GCaMP6s imaging analysis. For drug/metabolite cocktail, ***F***_***o***_ represented the average MFI of the first 10 frames. Area under the curve (AUC) quantification was performed using GraphPad Prism v10 software for each mouse.

Peak 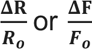 was calculated as the maximum for each ROI. Trials with excess z-movement (such that ROIs were no longer visible for >50% of the run) were eliminated and portions of trials in which ROIs were no longer visible were trimmed.

#### FUNCAT quantification

Images were imported into FIJI (ImageJ) for quantification. For FUNCAT % area quantification 20x tiled images containing somatosensory and motor cortex were z-projected (max projection), and ROIs were drawn around L2/3 and L5 cellular layers in both regions. FUNCAT signal background was subtracted using the “subtract background” (rolling ball radius 50 pixels), and a “median filter” was applied (pixel radius 2). Images were then thresholded identically and the “analyze particle” function was used to measure the thresholded FUNCAT area. Percent area was calculated as the proportion of FUNCAT positivity within the full layer ROI. 2-4 slices were used per mouse and averaged.

To calculate the FUNCAT signal within neurons generally (immunostained NeuN+ cells) or excitatory neurons (tdt+ cells from Camk2a^Cre/+^:R26^tdTomato/+^ mice), confocal 40x images were z-projected (maximum projection) and split. The NeuN or tdT channel was background subtracted (rolling ball radius 50 pixels) and median filtered (radius 2 pixels) and thresholded to include the maximum number of cells while reducing noise. After thresholding, the “watershed” function was used to separate touching cells and the “analyze particle” function (size: 5-infinity, 0.15-1 circularity) was used to generate neuron ROIs. MFI within each ROI was calculated. 2-4 slices were used per mouse. For fold change calculations, the average neuron MFI per section was calculated and represented 1 technical replicate. Technical replicates from the same layer, same mouse, same staining session, and imaged on the same day were averaged to get the average layer specific neuron MFI per mouse, which were considered biological replicates. Biological replicates were divided by the average of control group replicates to calculate the fold change per replicate (mouse).

To calculate the FUCNAT signal within microglia (immmunostained Iba1+ cells), a custom written FIJI macro was employed: Channels were split, and a median filter (1 pixel radius) was applied to the FUNCAT and IBA1 channels. Background from the IBA1 channel was subtracted (rolling ball radius 50 pixels). The IBA1 channel was thresholded identically for all images and converted to a mask. The new “mask” channel was combined with the FUNCAT channel using the “image calculator” “and” function to generate the microglia FUNCAT signal. All values outside of the mask were converted to value 0 and the resulting image was projected. ROIs were drawn manually for the full cell and soma. The fluorescent integrated density was measured in each ROI to calculate total FUNCAT signal per microglia and per microglia soma. Fold change calculations were calculated as with neuronal FUNCAT.

#### Microglia Morphology

Confocal image stacks from L2/3 and L5 were imported into Imaris 10.2 software (Oxford Instruments) for subsequent morphological analyses. In Imaris, surfaces representing entire IBA1-positive microglia were generated per image using the Machine Learning Segmentation, applying a smooth surface detail parameter of 0.155 µm. These surfaces were then utilized to create masked channels isolating microglial signals from the background. Soma-specific surfaces were subsequently segmented from these masked channels using Imaris’ Machine Learning Segmentation method tool. All somas generated via automatic segmentation were included in the analysis.

Filament tracing using the Machine Learning Segmentation method was performed on the masked channels to reconstruct microglial processes, with object-object statistics enabled to quantify morphological parameters comprehensively. After automatic filament generation, 5-10 microglia were randomly selected per image, manually inspected, and cleaned up to ensure accurate filament tracing before being included in the filament analysis. Creation parameters for surface and filament generation were saved and applied consistently across all images for batch analysis, ensuring uniformity and reproducibility of segmentation settings.

Morphometric data—including soma size, filament length, branching complexity, and other relevant parameters—were automatically extracted in Imaris 10.2. Data cleaning and organization were conducted in R Studio.

#### GLUT1 vessel quantification

Confocal image stacks covering motor cortex (immediately above dorsal peak of corpus callosum) were imported into FIJI (ImageJ). A custom written FIJI macro was used to calculate the percent of GLUT1 area within lectin (TL)+ blood vessels: The TL channel was z projected (max intensity), background subtracted (rolling ball=50), median filtered (radius=3), thresholded (identically for each section compared), converted to a mask, eroded 2x with the “erode function”, and dilated 2x with the “dilate” function. The mask was selected with “create selection”, which highlighted all TL+ vessels, and the total TL area was measured, which represented the total vessel area considered. The GLUT1 channel was selected and the “restore selection” was applied. Within the restored section the GLUT1 area was thresholded (identically for compared samples), converted to a mask, and measured, which represented the total GLUT1 area. The GLUT1 area was divided by the TL area (*100) to generate the “GLUT1% Vessel Area.” Fold change was measured as above.

### Flow Cytometry

WT mice randomized to either running or HC conditions. Running mice ran on a rotarod (as with *in vivo* FUCNAT) for 1 hour after which they were returned their home cage for 30 mins prior to brain isolation while HC mice remained in their HC until brain isolation. For brain immune cell isolation, to prevent artifactual transcriptional signatures induced by cell processing, mice were perfused with 20mL of an inhibitor cocktail of 5µg/mL actinomycin D (A9789, Sigma) and 10µM triptolide (T3652, Sigma) in PBS with 5mM EDTA^98^. All samples were kept on ice or at 4°C unless otherwise noted. Following perfusion, the cortex was dissected out, and enriched motor cortex (rostral of bregma to the olfactory bulb) was minced and digested with collagenase D 240U/ml (11088858001, Sigma) in PFH solution: 2% fatty acid free BSA (700-107P-100, Gemini Bio), 1mM hepes buffer in PBS, with 5ug/mL actinomycin D, 10µM triptolide, and anisomycin 27.1 µg/ml (A9789, Sigma) for 30 mins at 37°C. Following digestion, a cell slurry was made with a 3mL syringe and tissue was mashed through a 100µM cell strainer. The single cell suspension was centrifuged at 1500 rpm for 5 mins to pellet the cells. After centrifugation, the supernatant was aspirated, and the pellet was resuspended in 10ml 35% Percoll in PFH (p4937, Sigma) and spun at 2000RPM for 30 mins at RT with no break. After centrifugation, the supernatant containing myelin was aspirated, the cells were resuspended in cold PFH, filtered again, and centrifuged at 1500 rpm for 5 mins. For endothelial cell/astrocyte isolation, the motor cortex was processed using the Adult Brain Dissociation Kit (Miltenyi Biotec, 130-107-677) per manufacturer’s instructions using the 37C_ABDK_02 gentleMACS program and the gentleMACS Octo Dissociator. For brain immune cell staining the following cell-surface markers/viability dies were used, diluted in PBS: Live/Dead e780 (1:1000, 65-0865-14, ThermoFisher), Rat anti-CD45 BUV395 (1:200, 564729, BD), Rat anti-CD11b BV421 (1:400, 101251, BioLegend), Mouse anti-CD64 BV650 (1:200, 740622, BD), Rat anti-MHC2 BV711 (1:200, I-A/I-E, 107643 BioLegend), Rat anti-CD206 PE/CY7 (1:200, 141720, BioLegend) for 1hr at 4°C. For astrocyte/endothelial preps the following cell-surface markers/viability dyes were used: Live/Dead fixable Blue (1:500, L23105, Molecular Probes), Rat anti-CD31 PE/CY7 (1:100, 25-0311-82 Invitrogen), Rat anti-CD45 PE (1:100, 103106, BioLegend), mouse anti-GLAST APC (1:50, 130-123-555, Miltenyl Biotec). Samples were then washed, pelleted, and fixed/permeabilized using the Foxp3/transcription factor staining buffer set (00-5523-00, ThermoFisher) per manufacturer’s instructions. Samples were then blocked in 2% normal mouse serum (015-000-120, Jackson Immuno) for 15 min at RT and stained for intracellular protein Cyr61 using rabbit anti-Cyr61/CCN1 Alexa Fluor 488 (1:300, NB100-356AF488, Novus) for 30 mins at RT. Samples were then washed 2x with permeabilization buffer and resuspended in PBS before analyzing on a BD FACS Symphony A5.

Data was imported into FlowJo V10 (FlowJo LLC) for analysis. Microglia were gated Live/CD45^+^/CD11b^+^/CD64^+^/CD206^-^/MHC2^-^. CD206^High^ BAMs were gated as Live/CD45^+^/CD11b^+^/CD64^+^/CD206^+^/MHC2^-^. MHC2^+^ BAMs were gated as Live/CD45^+^/CD11b^+^/CD64^+^/^mid/low^/MHC2^+^ per^72^. Endothelial cells were gated as Live/CD45^-^

/CD31^+^. Astrocytes were gated as Live/CD45^-^/GLAST^+^. All samples were compared to Cyr61 fluorescence minus one controls to determine %Cyr61+ and normalized MFI.

For experiments using *in vivo* brefeldin A (BFA)^75^, running or HC mice were injected retro-orbitally with .25mg of BFA each (reconstituted to 20mg/mL in DMSO and further diluted in PBS) 2.5 hours before training or HC resting. All mice were perfused 4 hours after BFA injection.

### TRAP-Seq (All steps were performed in RNAse-free conditions)

#### Bead preparation

Streptavidin-conjugated magnetic beads were rinsed with 1x PBS and incubated with 120 mg of protein L conjugated to biotin for 35 min/rotating/room temperature (RT). Beads were washed 5x with PBS + 0.05% IgG and protease-free BSA and incubated with 50mg of each anti-GFP antibody (Htz-19C8 and Htz-19F7, Memorial Sloan-Kettering) for 1h/rotating/RT. Finally, beads were washed 3x with low-salt buffer (80 mM HEPES KOH pH 7.3, 600 mM KCl, 40 mM MgCl2, 1% NP40, 0.5 mM DTT, protease/phosphatase inhibitors and 100 mg/ml Cycloheximide). Beads were resuspended in low-salt buffer and stored for a maximum of 24h at 4°C prior to experiment.

#### Animal training and sample collection

Cx3cr1^CreERT2/+^:R26^EGFP-L10a/+^ mice that had been gavaged with tamoxifen at least 3 weeks prior to experimentation were either trained on a rotarod for 1 hr (See “Rotarod *in vivo* FUNCAT”) and returned to their home cage for 30 mins or left in their home cage for the duration of the experiment. Following training or home cage resting, enriched motor cortex was collected in ice-cold PBS supplemented with 100 mg/ml Cycloheximide and immediately lysed in an ice-cold glass pestle container with 1 ml of tissue lysis buffer (80 mM HEPES KOH pH 7.3, 600 mM KCl, 40 mM MgCl2, 1% NP40, 0.5 mM DTT, protease/phosphatase inhibitors, 300U RNAsin and 100 mg/ml Cycloheximide). Tissue was homogenized with 12 strokes of a motorized pestle and suspension transferred to an Eppendorf tube. Homogenate was cleared by centrifugation (2000 *g*/ 10 min./4°C). Supernatant was saved and supplemented with 1/9^th^ v:v of 300 mM DHPC, mixed by inversion, incubated for 5 min in ice, and centrifuged at 20.000 *g*/10min/4oC. 1/10^th^ of the volume was saved for total input RNA-sequencing, and the remaining was added with 200 ml of magnetic beads and incubated at 4°C/rotating/overnight. Samples were then briefly spun to recover all beads and washed 4x with 1 ml ice-cold high-salt buffer (80 mM HEPES KOH pH 7.3, 1.4 M KCl, 40 mM MgCl2, 1% NP40, 0.5 mM DTT, protease/phosphatase inhibitors, 300 U RNAsin and 100 mg/ml Cycloheximide). Beads were then resuspended in 350 ml room temperature RLT buffer added with 10% b-mercaptoethanol (RNEasy plus kit, Qiagen). 350 ml of RLT buffer + b-mercaptoethanol was added to input samples as well. RNA purification was done using RNEasy plus kit, following standard procedures suggested by the manufacturer. RNA quality was measured using Agilent’s ScreenTape for global assessment of RNA integrity.

#### Library preparation and sequencing

The SMART-Seq HT Kit (Takara) was utilized to generate complementary DNA (cDNA) from 1 ng of total RNA input. The synthesis process included 13 PCR cycles. The quality and concentration of the stock RNA were assessed using an Agilent Pico Chip on the Bioanalyzer system, ensuring high-quality RNA inputs. cDNA was quantified using the Invitrogen Quant-iT system and subsequently diluted to a concentration of 0.3 ng/µL for library preparation. Libraries were prepared using the Nextera XT DNA Library Preparation Kit, employing a total of 12 PCR cycles to amplify and uniquely barcode the samples. The quality of the generated libraries was assessed using the Agilent High Sensitivity TapeStation to verify fragment sizes and integrity. Additionally, the concentration of libraries was measured with the Invitrogen Quant-iT system to ensure accurate normalization prior to pooling. Libraries were pooled in equimolar ratios and sequenced on the Illumina NovaSeq X+ platform using paired-end 100-cycle reads to a depth of 50 million reads per sample. Fastq files integrity was tested using md5 checksum and FastQC pipeline. The sequencing data was processed using standard bioinformatics pipelines.

#### Alignment to the genome and feature count

Sequencing adapters and low-quality bases were trimmed using *Cutadapt* version 1.8.2 using the following arguments (-q 20 −O 1 -a CTGTCTCTTATA). Trimmed reads were then aligned to the mm10 mouse genome assembly using STAR version 2.7.7a and raw gene counts were obtained using *-quantmode* argument^99^. Total feature counts were obtained using *featureCounts*^100^. Feature counts were extracted and converted into one single raw count matrix, where each row was a gene, and each column was a sample. When samples were obtained from different cohorts of mice, we used ComBat-Seq to batch-correct^101^.

#### Differential expression analysis and Gene Set Enrichment Analysis (GSEA)

Identification of DEGs was made using DESeq2 package, from R^102^. Briefly, crude count matrices were batch corrected using ComBat-seq^101^. The resulting matrix served as input to DEseq2. Transcripts with less than 20 counts in more than half of the samples were filtered out. Datasets were then normalized by the size of library, and variance stabilized using the *vst*() function from DEseq2 package. DEG identification was done by filtering genes that had *p* adjusted value < 0.05 in comparisons between conditions. A complete list of all DEGs from all datasets is available in Supplementary table 3. These lists of DEGs were independently subjected to Gene Ontology analysis using the *GProfiler* package from R, with *P* value adjusted using false discovery rate. Heatmaps were generated using *ComplexHeatmap* package. Gene Ontology plots were done using the *ggplot2* package. To plot signed *p* adjusted plots, *p* adjusted of downregulated GOs were multiplied by −1 prior to plotting. GSEA was performed using the Broad Institute’s GSEA tool to assess the enrichment after generating a ranked list of genes based on Log_2_FC of Running vs HC transcripts and assessing enrichment of GO gene sets.

### Single Nucleus RNA Sequencing

#### Tissue preparation

Cx3cr1^CreERT2/+^:R26^DTR/+^ and Cx3cr1^CreERT2/+^:R26^+/+^ mice aged at 4-5-week-old were gavaged with tamoxifen as described above. Twenty-seven days after tamoxifen gavage, mice were IP injected DT for three consecutive days as above. One day after the last injection of DT, mice were anesthetized and decapitated. Fresh brain samples were quickly separated from the brain skull and split into two parts through the midline. The right brain hemisphere was cut with a blade to isolate the motor cortex and immediately frozen in liquid nitrogen for single-nucleus sequencing. All experimental protocols were approved by the Institutional Animal Care and Use Committee of Shenzhen Graduate School, Peking University (approved protocol 11514), and adhered to institutional guidelines.

#### Single-nucleus suspension preparation

Tissue samples were subjected to nuclei isolation prior to library construction. Flash-frozen tissues (∼100 mg) were homogenized in 2 mL ice-cold Homogenization Buffer (250 mM sucrose (Ambion), 10 mg ml–1 BSA (Ambion), 5 mM MgCl2 (Ambion), 0.12 U μl–1 RNasin Plus (Promega, N2115), 0.12 U μl–1 RNasein (Promega, N2115) and 1× Complete Protease Inhibitor Cocktail (Roche, 11697498001)) using a pre-chilled tissue grinder. After 3-5 min incubation, tissues were manually ground until resistance diminished. The homogenate was filtered sequentially through 70-µm and 40-µm strainers, and the filtrate was collected. Nuclei were pelleted (500 ×g, 5 min, 4°C), washed twice with 1 mL Wash Buffer (Homogenization Buffer containing 1% Igepal (Sigma, CA630)), and resuspended in Blocking Buffer (320 mM sucrose, 10 mg ml–1 BSA, 3 mM CaCl2, 2 mM magnesium acetate, 0.1 mM EDTA, 10 mM Tris-HCl, 1 mM DTT, 1× complete Protease Inhibitor Cocktail and 0.12 U μl–1 RNasein). Nuclei were counted directly if the suspension was clean; otherwise, fluorescence-activated sorting (FACS) was performed to remove debris.

Single-nucleus suspensions were processed following the DNBelab C Series Single-Cell Library Prep Set (MGI, 1000021082) workflow. Briefly, single nuclei were subjected to droplet encapsulation, emulsion breakage, bead collection, reverse transcription, cDNA amplification, and purification to generate barcoded cDNA libraries. Indexed sequencing libraries were constructed according to the manufacturer’s protocol. Library concentrations were quantified using the Qubit ssDNA Assay Kit (Thermo Fisher Scientific, Q10212). Sequencing was performed on a DNBSEQ platform at the China National GeneBank (Shenzhen, China).

#### Raw data processing

Raw FASTQ reads were filtered and demultiplexed using PISA (https://github.com/shiquan/PISA), then aligned to the mouse reference genome (mm10) using STAR. To account for unspliced transcripts in nuclei, a custom ‘pre-mRNA’ reference was generated for alignment of count reads to introns as well as to exons. A nucleus versus gene UMI count matrix was constructed with PISA.

#### Dimensionality reduction, clustering, and cell type identification

We applied the following preprocessing and quality-control filters to each nucleus: a minimum of 800 detected genes, total UMIs ranging from 1,400 to 20,000, and inclusion of only genes expressed in at least three nuclei. Nuclei with more than 5% mitochondrial gene counts and 10% ribosomal gene counts were removed. Potential doublets were identified and excluded using the R package DoubletFinder. Dimensionality reduction and clustering were performed using the Seurat workflow (https://github.com/satijalab/seurat). Briefly, the gene expression matrix was normalized and scaled, subsequent principal component analysis. The optimal number of principal components was chosen by using the ElbowPlot, for clustering with the specific resolution parameters. Using R package Harmony for batch correction.

Clusters were annotated using canonical markers of known cell types in combination with the distinct marker signatures identified. For each cell type, we used cell type specific/enriched marker genes to determine cell type identity. Prior to downstream analysis, identified clusters (Fig. 1E) were combined to the following groups: excitatory neurons (ExN; including L2/3 IT, L4/5 IT, L5 IT, IT, L6 IT, L5 PT, L6 CT, L6b, and L5/6 NP), inhibitory neurons (InN; Pvalb, Sst, Vip, and Lamp5), astrocytes (Ast; Astrocyte 1-3), oligodendrocytes (OLG; OLG 1-2), vascular cells (Vas; Endothelial, VLMC, and Ependymocytes). Neuronal clusters (Neu) comprised both ExN and InN populations.

#### Gene differential expression and pathway analysis

Differentially expressed genes (DEGs) were calculated using the FindMarkers or FindAllMarkers function in Seurat (Wilcoxon rank-sum test). Protein-coding genes with absolute value average log2FC > 0.25, FDR-adjusted P < 0.01, expressed in at least 20% of cells, were considered significant. Gene-set enrichment analysis (GSEA) was performed using the Broad Institute’s GSEA tool to assess the enrichment of Hallmark gene sets and Gene Ontology Biological Process (GOBP) gene sets (FDR < 0.1).

### Statistical Analysis

Statistical analysis and data visualization were performed in GraphPad Prism (v10) or R studio. Details regarding tests used and number of replicates for each experiment are described in figure legends. Statistical significance was determined by two-tailed unpaired Student’s t-test when comparing two independent groups or paired t-test when comparing values from the same group. Wilcoxen signed-rank test was used in paired sequencing data when normality could not be assumed, grouped data were analyzed by two-way ANOVA or a mixed-effects model if data contained repeated measures. Individual group differences were determined by Tukey’s post hoc test for two-way ANOVAs or Sidak’s post hoc test for repeated measures mixed-effects model with α set to .05. Significant interactions are reported in figure legends. All data are presented as individual values and mean ± S.E.M or truncated violin plots with maximum values, quartile ranges, and median value. Simple linear regression was used to assess correlation.

## Code Availability

Any additional information required to reanalyze the data reported in this paper is available from the lead contact upon request.

---

**Figure S1.**
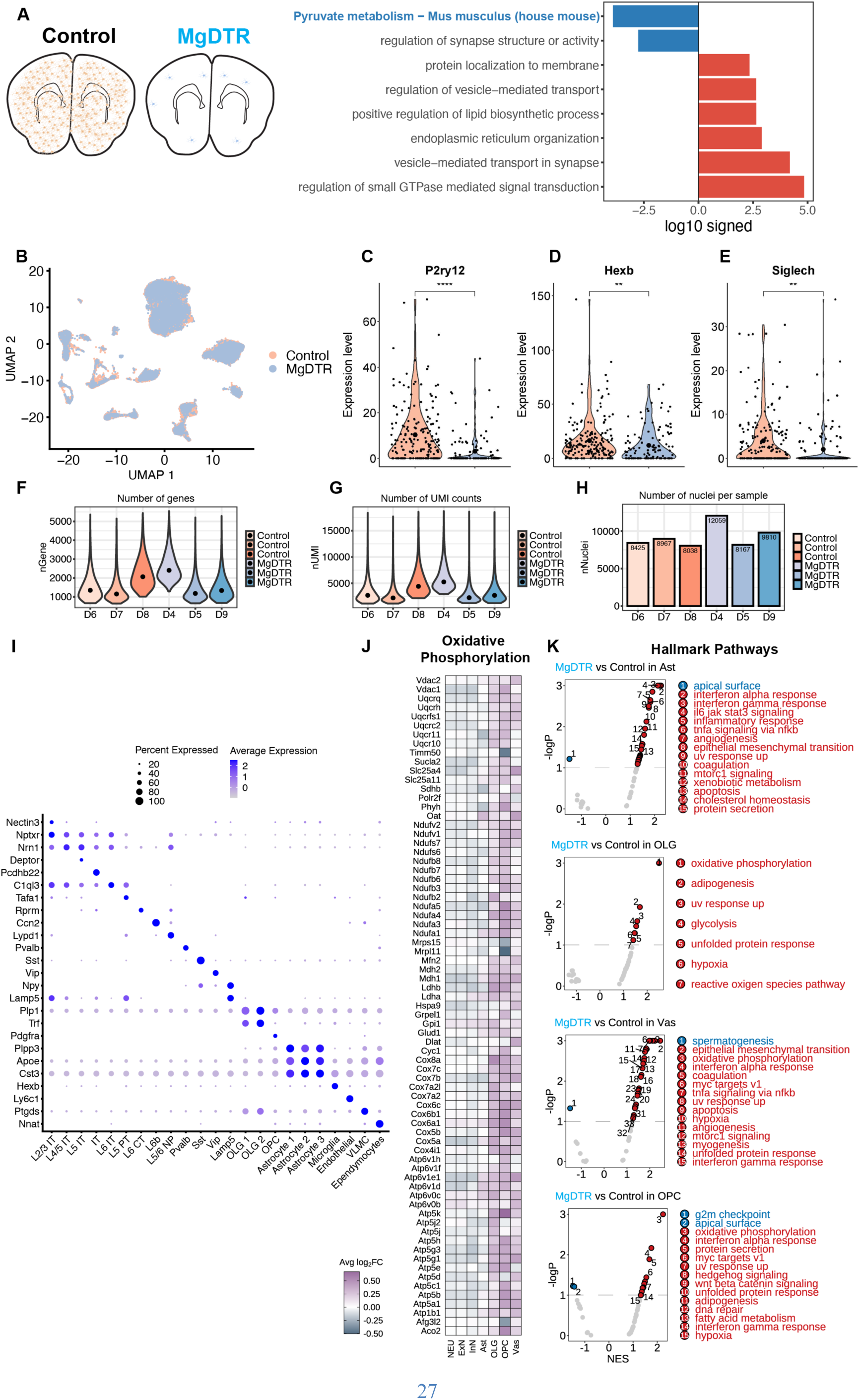
Multi-omic evaluation of motor cortical changes after microglia depletion, related to Figure 1. (**A**) Re-analysis of top dysregulated protein sets (Blue=down in MgDTR, Red=up) after shotgun proteomics comparing full brains of Control vs. MgDTR mice from *Parkhurst et al., 2013*^16^ using Metascape^94^. (**B**) UMAP depicting the relative distribution nuclei in each cluster from Fig. 1C between control (orange) and MgDTR (blue) groups. Relative expression levels of canonical microglia genes (**C**) P2ry12, (**D**) Hexb, and (**E**) Siglech between nuclei in control and MgDTR groups. (**F**) Number of genes captured, (**G**) unique molecular identifiers (UMI), and (**H**) nuclei per replicate (N_control_= 3 mice and N_MgDTR_=3 mice. (**I**) Heatmap depicting canonical markers (y axis) used to identify clusters (x axis). Dot size represents percent of nuclei within the cluster that expressed the y axis marker and shade of blue represents average expression per nuclei. (**J**) Fold change heatmap for all genes within the GOBP “Oxidative Phosphorylation” pathway between MgDTR and Control mice for aggregate clusters using GSEA. (**K**) Top 15 GSEA “Hallmark Pathways” discovered with FDR q<.1 within astrocytes (Ast), oligodendrocytes (OLG), vascular cells (Vas), and OPCs (see “*Dimensionality reduction, clustering, and cell type identification”* in materials and methods for aggregate cluster definitions). Wilcoxen signed-rank test for (C), (D), and (E). **P<.01, ****P<.0001. GOBP=Gene Ontology Biological Pathways, GSEA= Gene Set Enrichment Analysis.

**Figure S2.**
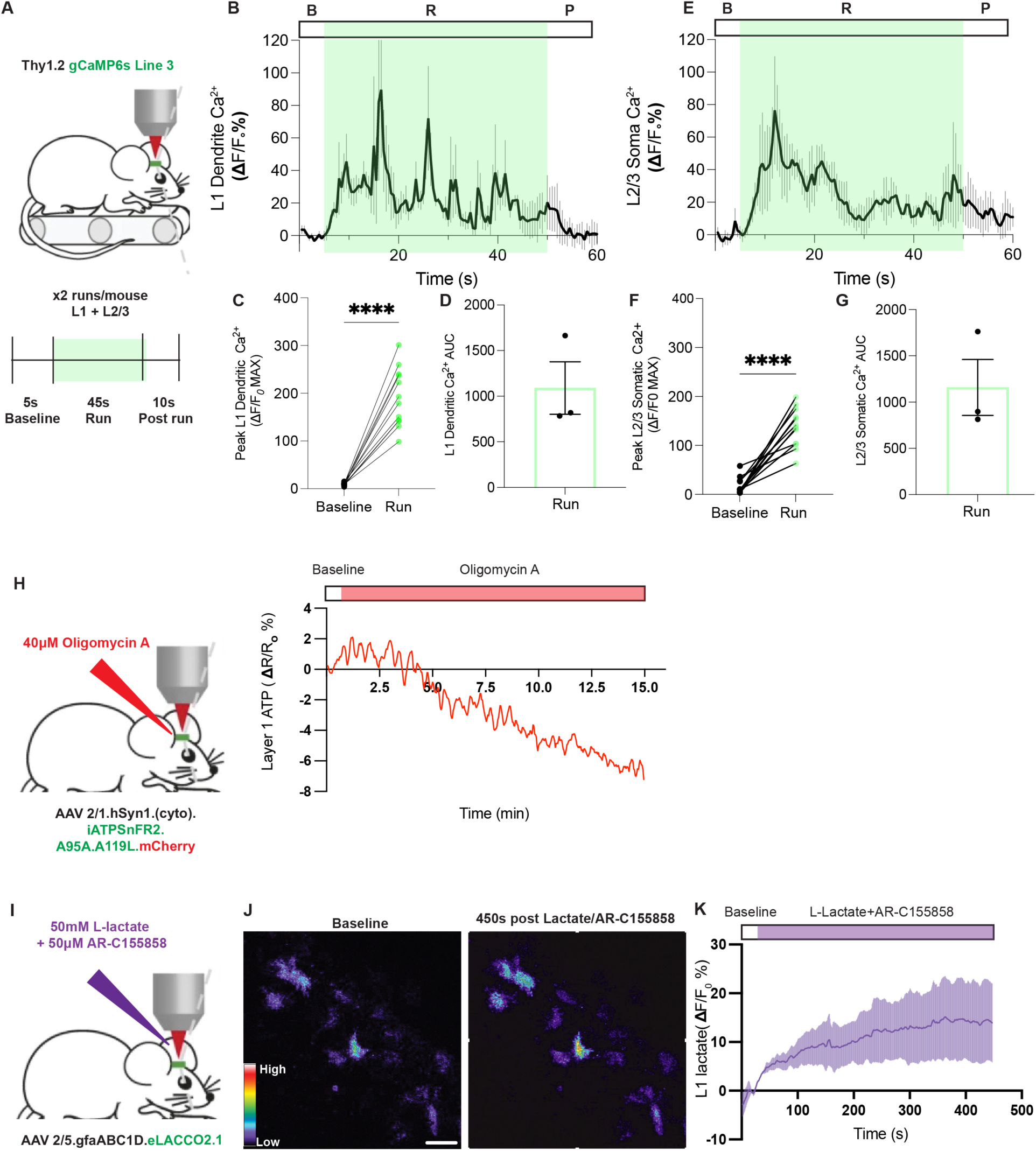
Validation of motor cortical neuronal activity and metabolic sensor specificity *in vivo*, related to Figure 2. (**A**) Schematic illustration of experimental paradigm for in vivo neuronal calcium imaging in Thy1.2 gCaMP6s line 3 mice which express gCaMP6s in neurons. Motor cortical neuronal somas and dendrites were imaged in L2/3 and L1 during 2 45s runs. (**B**) Mean 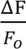 trace per mouse during baseline, run, and post run states (N=3 mice, average of 4 dendrites per mouse). (**C**) Peak 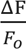 per dendrite during baseline and run phases (N=12 dendrites). (**D**) Run phase integrated Ca^2+^ per mouse (N=3 mice). (**E** to **G**) Similar to (B to C) but for L2/3 somas (N=12 somas, N=3 mice). (**H**) Schematic illustration and trace of L1 neuropil iATPSnFR2 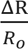 after application of oligomycin A to an open cranial window normalized to a 10-frame baseline (N=1 mouse). (**I**) Schematic illustration of experiment in which 50mM of L-lactate and 50µM MCT inhibitor AR-C155858 were added directly to an open cranial window and imaged for 450s. (**J**) representative image of astrocytic eLACCO2.1 fluorescence at the beginning and after 450s of lactate/drug cocktail. (**K**) Mean 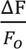 eLACCO2.1 trace per mouse normalized to first 10 frames (N=2 mice). Two-tailed, paired *t* test for (C) and (F). ****P<.0001. Scale bar: 50 µm for (J). Shading represents +/- S.E.M.

**Figure S3.**
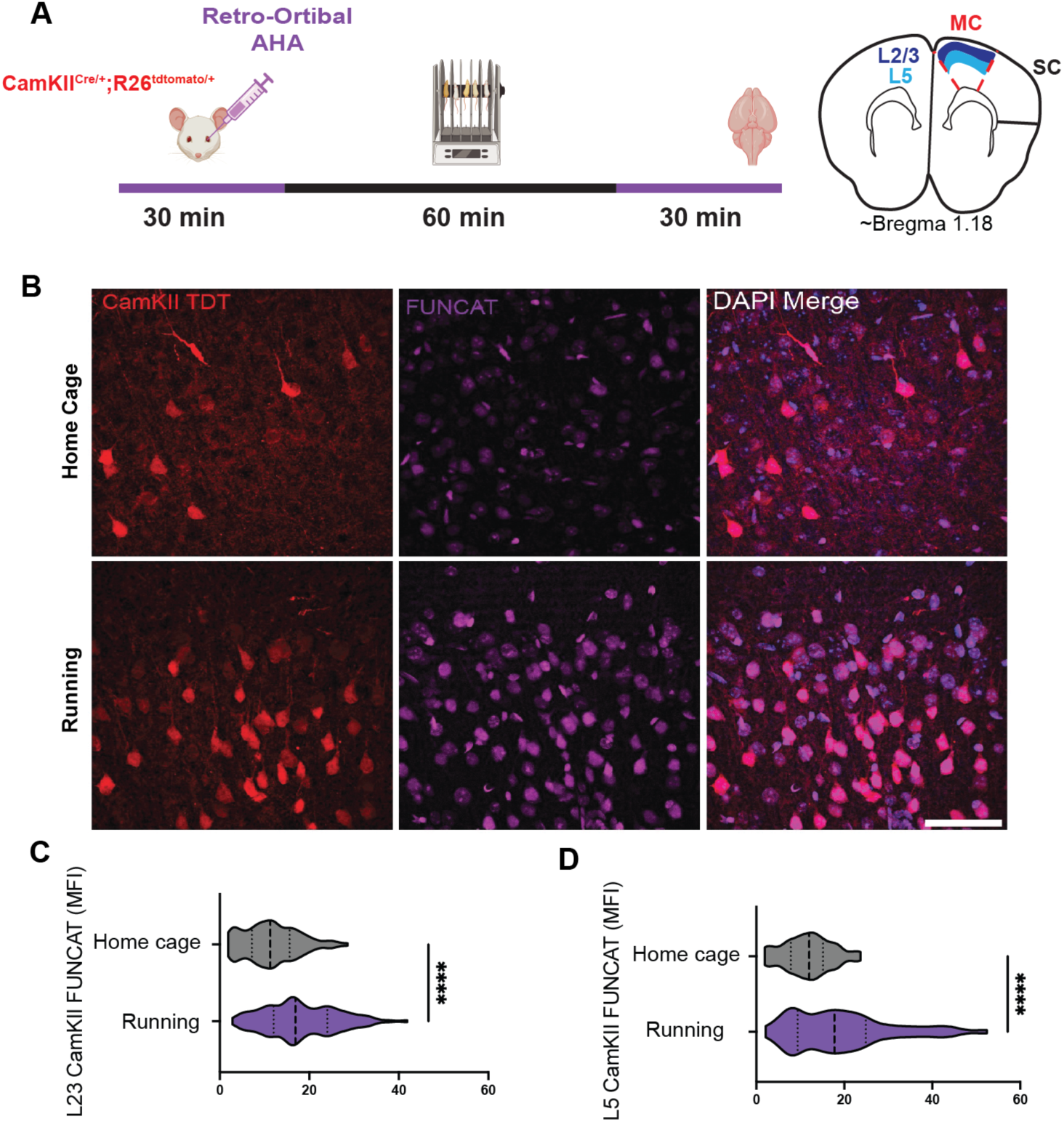
Running leads to increased protein synthesis in motor cortical excitatory neurons, related to Figure 3. (**A**) Schematic illustration of motor training *in vivo* FUCNAT experiment in CamKII^CRE/+^;R26^tdtomato/+^ mice which express fluorescent protein tdtomato in CamKII expressing excitatory neurons. (**B**) Representative images of L2/3 excitatory neurons (red) and FUNCAT labeling (purple) from HC and running mice. Quantification of excitatory neuron FUNCAT mean fluorescence intensity (MFI) in (**C**) L2/3 and (**D**) L5 (N_HC2/3_=101 cells, N_Run2/3_=253 cells, N_HC5_=47 cells, N_Run5_=124 cells from 2 mice per condition). Two-tailed, unpaired *t* test for (C) and (D). ****P<.0001. MC= motor cortex, SC= somatosensory cortex. Scale bar: 50 µm for (B). Illustration (A, left) created with BioRender.com.

**Figure S4.**
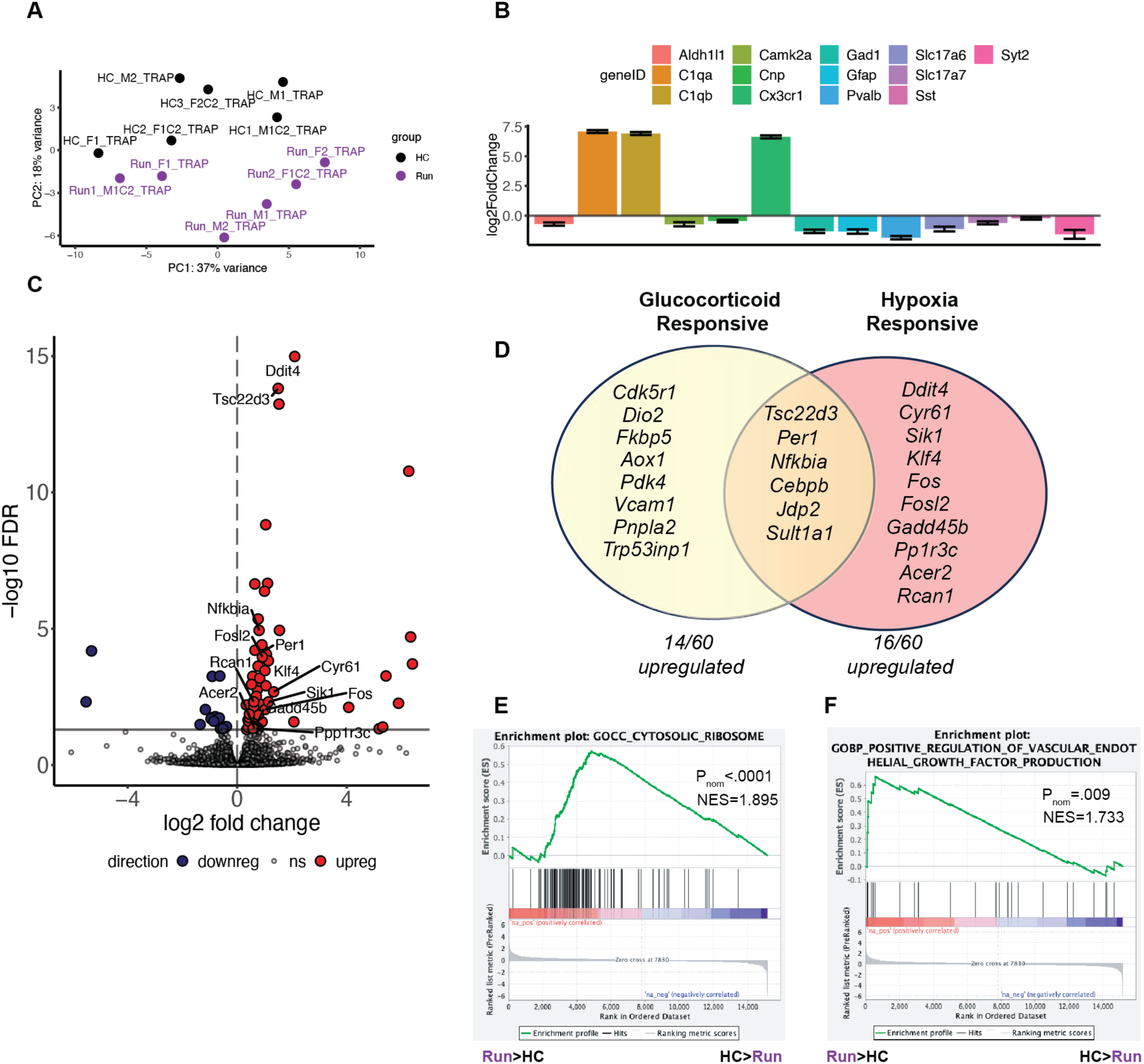
Microglia TRAP-seq reveals distinct transcriptional changes during running, related to Figure 5. (**A**) Principal component analysis of HC (black) and running (purple) replicates from TRAP-seq experiment (N=6 mice per group). (**B)** Bar plot depicting log_2_FC of cell-type specific transcripts in TRAP bound fraction/unbound total lysate fraction from TRAP-seq experiment. (**C**) Volcano plot with upregulated (red, 60 total) downregulated (blue, 17 total), and non-significant (grey, P_adjusted_>.05) mRNAs in microglia from running mice. (**D**) Ven diagram of genes identified as glucocorticoid (yellow circle), hypoxia responsive (orange), or both. Genes were identified using compiled glucocorticoid responsive and hypoxia responsive gene sets from (58). Beneath is the number of genes within each category out of the total upregulated genes. (**E**) GSEA reveals the over-representation of genes involved in in mRNA translation such as “cytosolic Ribosome” and (**F**) genes involved in the “Positive Regulation of Vascular Endothelial Growth Factor Production” in microglia from running mice. The left side of each plot represents transcripts ranked higher in microglia from running mice and the right side represents transcripts ranked higher in HC mice. (E) ****P_nominal_<.0001, (F) **P_nominal_=.001. NES= Normalized Enrichment Score.

**Figure S5.**
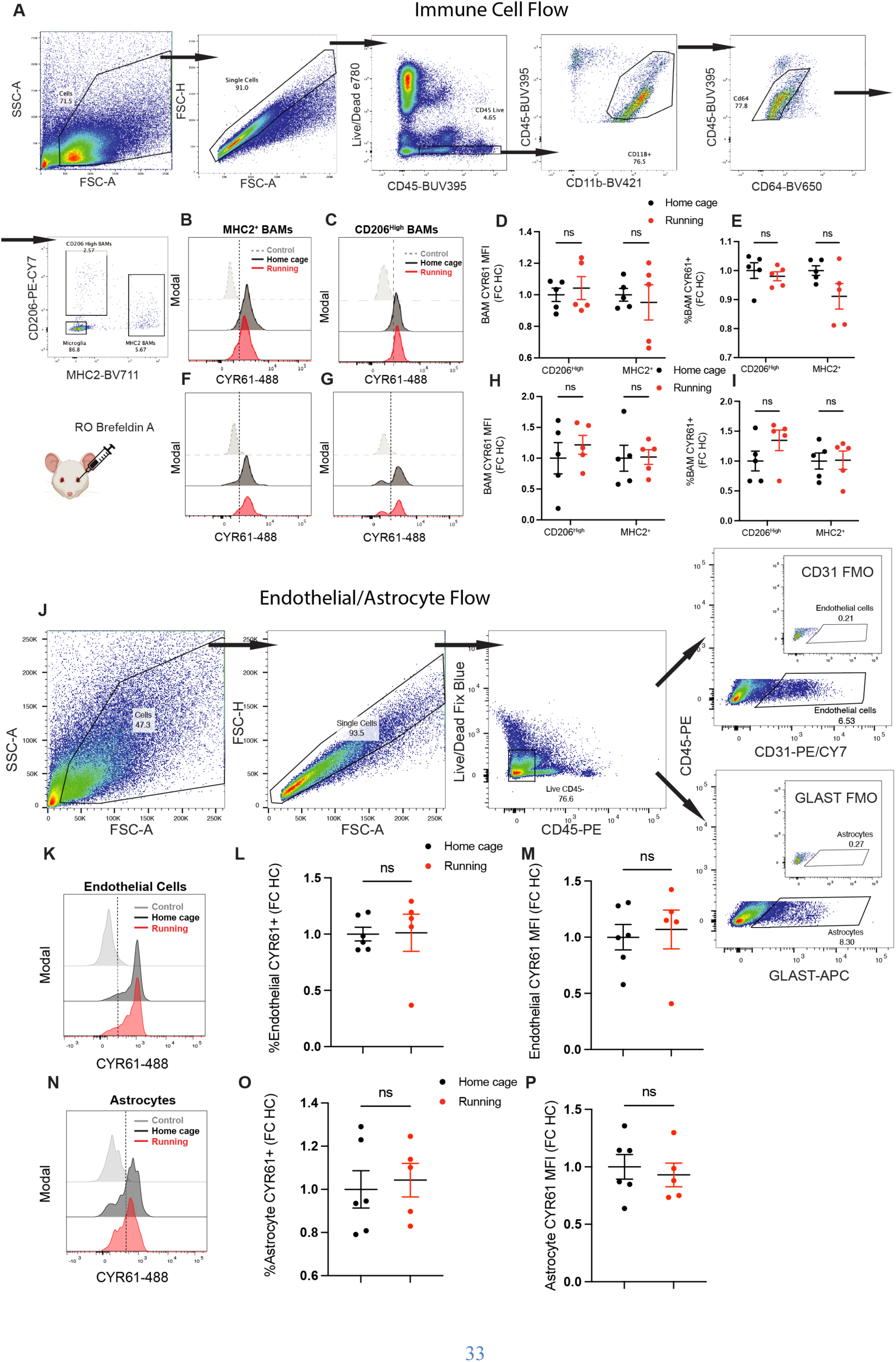
Running does not change intracellular CYR61 protein levels in BAMs, endothelial cells, or astrocytes, related to Figure 6. (**A**) Gating strategy for identification of microglia, CD206^high^, and MHC2^+^ BAM populations. Representative histograms depicting CYR61 positivity (to the right of dotted line) in HC (black) and Running mice (red) compared to FMO control (grey) within (**B**) MHC2^+^ or (**C**) CD206^high^ BAMS. Quantification of CYR61-488 (**D**) fold change (FC) mean fluorescent intensity (MFI) and (**E**) percent positivity per mouse for CD206^high^ and MHC2^+^ BAM populations from HC (black dots) or Running (red dots) mice (N_HC_=5, N_Run_=5) normalized to HC mean for each cell type. (**F-I**) Similar to (B to E) but for an experiment in which brefeldin A was injected 2.5 hours prior to motor training (see Figure 7D). (**J**) Gating strategy for identification of endothelial cells and astrocytes with respective FMOs (insets) used for gating. (**K-M**) Similar to (C-E) but for gated endothelial cells. (**N-O**) Similar to (C-E) but for gated astrocytes. (L, M, O, and P) Unpaired two-tailed student’s t-test. (D, E, H, and I) Repeated measures mixed effects model with Sidak’s post hoc test, ns= not significant. FMO= Fluorescence Minus one controls. Illustrations created with BioRender.com.

